# Genome-wide Transcriptomic Analysis of T*oxoplasma gondii* Reveals Stage-specific Regulatory Programs and Metabolic Adaptations Driving Oocyst Sporulation

**DOI:** 10.1101/2025.09.10.674405

**Authors:** Electine Magoye, Jessica J. Bader, Chandra Ramakrishnan, Adrian B. Hehl, Marc U. Schmid

**Affiliations:** Institute of Parasitology, University of Zurich, Winterthurerstrasse 266a, Zürich, 8057, Switzerland; MWSchmid GmbH, Glarus, Switzerland

## Abstract

The sporulation of *Toxoplasma gondii* oocysts is a critical developmental transition transforming non-infective environmental stages into highly resilient and infective forms capable of zoonotic transmission. Despite significant advances, the molecular regulation underlying this complex process remains poorly understood. We generated comprehensive RNA-Seq datasets capturing gene expression dynamics at three timepoints during oocyst sporulation, to illuminate essential regulatory mechanisms and metabolic adaptations. Employing stringent variance filtering, unsupervised clustering, and differential expression analyses, we identified transcriptional patterns associated with key developmental transitions, notably the involvement of *T. gondii* Apetala2 (TgAP2) transcription factors and other DNA-binding proteins. We highlight significant shifts in metabolic pathways essential for oocyst environmental resilience and infectivity, including lipid metabolism, amino acid biosynthesis, and specialized secondary metabolism. Furthermore, we investigated expression profiles of genes associated with structural development of oocyst and sporocyst walls, environmental persistence, and host transmission. Importantly, combining these datasets with prior life cycle datasets covering intermediate and final host stages, we close an important knowledge gap by providing a genome-wide mRNA expression profile spanning the entire *T. gondii* life cycle. Integration with previous foundational transcriptomic and proteomic work provides refined resolution and identifies novel candidate genes and pathways. Collectively, these findings substantially advance our understanding of *T. gondii* sporulation in the wider context of the development in the cat intestine, highlighting critical molecular events underpinning parasite transmission and environmental adaptation.

## Background

The sporulation of *Toxoplasma gondii* oocysts is a complex and critical stage in the parasite’s lifecycle, transitioning from an immature, unsporulated and thus non-infective state into a sporulated form capable of infecting a broad range of intermediate hosts, including humans. *T. gondii* is a widespread apicomplexan parasite, closely related to other medically significant pathogens such as *Plasmodium spp*. and *Cryptosporidium spp*. It can infect all warm-blooded animals, highlighting its remarkable adaptability and zoonotic potential. Oocysts are highly infectious for humans and livestock, causing severe congenital toxoplasmosis with miscarriages or neurological deficits, blindness, and other significant developmental disorders in newborns and abortions in small ruminants.

Due to its capacity to infect multiple host species, *T. gondii* has become a prominent model organism for studying host-parasite interactions and immune evasion strategies. Tachyzoite stage parasites can be cultured in human and animal cells and induction of cysts and differentiation to bradyzoites makes *T. gondii* a great model to study *in vitro*. However, despite some recent advances, the development of pre-sexual and sexual enteroepithelial stages (EES) of the parasite in vitro remains to be established.

A significant public health concern arises from environmental contamination with sporulated oocysts, which enter the food chain and subsequently infect farm animals [1,2] and humans. Such zoonotic transmission poses considerable health risks, particularly in regions with intensive livestock production and inadequate sanitation. Despite the ubiquitous presence and environmental resilience of sporulated oocysts, the molecular processes governing sporulation remain poorly understood. Early foundational transcriptomic and proteomic analyses in *T. gondii* and *Eimeria tenella* have provided crucial initial insights into this stage, highlighting significant shifts in gene expression profiles and the appearance of proteins essential for environmental resistance and host cell invasion. Notably, pioneering work by Fritz et al. provided crucial insights into the changing transcriptome landscape during sporulation, identifying novel stage-specific genes and regulatory elements [3].

However, with respect to the precise molecular mechanisms for gene expression regulation during sporulation our understanding remains limited. This is especially true for the role of key transcription factors such as many members of the Apetala2 (AP2) family, as well as the metabolic shifts facilitating the formation of infectious oocysts and sporocysts. Moreover, detailed functional characterization of proteins involved in oocyst wall resilience and sporocyst formation is scarce.

Here we generated genome-wide RNA-Seq data of sporulating oocysts with the aim to capture gene expression patterns defining this key development in the external/environmental part of the *T. gondii* life cycle. Building upon previous datasets, including comprehensive transcriptional profiles of other lifecycle stages such as merozoites and gametocyte stages [4,5], this analysis with higher resolution focusing on early events during *T. gondii* sporulation when excreted oocysts experience changes in temperature and oxygen tension. Importantly, we close the gap between oocyst shedding into the environment and proliferating tachyzoites in the newly infected host allowing generation of a transcriptomic atlas of the full *T. gondii* life cycle as a basis for computational prediction of gene expression patterns with mechanistic implications. We identify and characterize genes specifically regulated by AP2 and other DNA-binding proteins, elucidate significant metabolic shifts occurring during sporulation, and define key proteins with roles in oocyst and sporocyst biogenesis. Collectively, this resource significantly narrows the existing knowledge gap in the transcriptional landscape of the *T. gondii* life cycle, offering critical molecular insights and highlighting novel candidates for targeted research and interventions in toxoplasmosis management

## Materials and Methods

### Ethics Statement

*T. gondii* strain CZ [6,7] clone H3 oocyst were harvested from experimental infections of cats (control groups) for a vaccine trial under the permit Nr. ZH040/17 of the veterinary Office of the Canton of Zurich (Switzerland) for which ethical approval from the animal rights protection commission of the Canton of Zurich (Switzerland) was required.

### Oocyst Isolation and Enrichment

Faeces from kittens infected with *T. gondii* CZ clone H3 collected at 5 – 6 days post infection were soaked in 0.1% Tween-80 in tap water and kept at 4 °C. The slurry was then sieved, washed and centrifuged at 1000 g for 10 min at RT. The sediments were then re-suspended in sucrose (55% w/w in tap water) and centrifuged at 1600 g, 10 min, 4 °C without breaks for deceleration in a swinging bucket centrifuge. Some tap water was dripped to the surface, stirred and the aqueous phase was transferred to a fresh tube. The tube was filled with cold tap water and spun at 1600 g as described above. Washing was repeated twice with water and once with Tris-EDTA buffer (50 mM Tris-Cl, 10 mM EDTA, pH 7.2-7.4); after last wash, leave small amount of supernatant. Overlay 1.15 g/ml CsCl in TE buffer with 1.10 g/ml CsCl and the upper layer with CsCl 1.05 g/ml (1.5 ml each for 1 ml oocyst slurry). The oocysts were then centrifuged at 16’000 g for 1 h at 4 °C. The top fractions including most of the CsCl 1.10 g/ml were place into a new tube and washed with 0.1% Tween-80/distilled H_2_O at 2500 g for 15 min at 4 °C. After removal of the supernatant, the oocysts were washed once in distilled water as above. Oocysts from several tubes were pooled and divided to 3 tubes and centrifuged at 2500 g as above. Oocysts from 1 pool were re-suspended in QIAzol Lysis Reagent (QIAGEN) and frozen at −80 °C.

### Sporulation

Oocysts from above were re-suspended in 2% H_2_SO_4_ and transfer to Petri dish. After 8 h or 8 d, oocysts were re-transferred to a tube and filled with distilled H_2_O. Oocysts were centrifuged at 2500 g for 15 min at 4 °C. The supernatant was removed and the wash repeated twice. Pellets were re-suspended in QIAzol Lysis Reagent and frozen at −80 °C.

### RNA Extraction

To extract RNA, aliquots of 600 µl were transferred to bashing bead tubes (Zymo ZR BashingBeads 0.1 mm and 0.5 mm). Oocysts were lysed in pre-cooled racks using a TissueLyser II system (QIAGEN) for 30 cycles/s and immediately placed in an ice/water bath. This procedure was repeated 4 times. The lysate was warmed to room temperature for 15-20 min, then centrifuged at 4750 rpm for 1 min. The supernatant was transferred to a fresh tube. An equivalent volume of 100% ethanol was added. The RNA was purified using the Direct-zol RNA Miniprep kit (Zymo Research) with the manufacturer’s protocol. An on-column DNase I digestion was performed for 15 min at RT after the first wash of the loaded columns.

### Library Preparation and RNA Sequencing

The quality of the isolated RNA was determined with a Qubit® (1.0) Fluorometer (Life Technologies, California, USA) and a Bioanalyzer 2100 (Agilent, Waldbronn, Germany). Only those samples with a 260 nm/280 nm ratio between 1.8–2.1 and a 28S/18S ratio within 1.5–2 were further processed. The TruSeq RNA Sample Prep Kit v2 (Illumina, Inc, California, USA) was used in the succeeding steps. Briefly, total RNA samples (100-1000 ng) were poly A enriched and then reverse-transcribed into double-stranded cDNA. The cDNA samples were fragmented, end-repaired and polyadenylated before ligation of TruSeq adapters containing the index for multiplexing Fragments containing TruSeq adapters on both ends were selectively enriched with PCR. The quality and quantity of the enriched libraries were validated using Qubit® (1.0) Fluorometer and the Caliper GX LabChip® GX (Caliper Life Sciences, Inc., USA). The product is a smear with an average fragment size of approximately 260 bp. The libraries were normalized to 10nM in Tris-Cl 10 mM, pH8.5 with 0.1% Tween 20.

The TruSeq SR Cluster Kit HS4000 (Illumina, Inc, California, USA) was used for cluster generation using 10 pM of pooled normalized libraries on the cBOT. Sequencing was performed on the Illumina HiSeq 4000 single end 100 bp using the TruSeq SBS Kit HS4000 (Illumina, Inc, California, USA).

### Normalization and Differential Expression Analysis

Adaptor sequences and low-quality stretches within the reads were removed with fastp (version 0.20.0 [8]). Transcripts were quantified with Salmon (version 0.12.0, options –type quasi-k 31 for the index build step and options –validate [9]). Mappings for the quantification step using the cDNA and gene annotation available from ToxoDB (*T. gondii* ME49, initially release 41, later release 68). Variation in gene expression was analysed with a general linear model in R with the package DESeq2 (version 1.20.0 and 1.24.0 [10]) according to a design with a single factor comprising all time points. Specific conditions were compared with linear contrasts. Specifically, all time points were compared to each other and the latest time point was compared to the average of the two earlier time points. Within each comparison, *p-*values were adjusted for multiple testing (Benjamini-Hochberg) and genes with an adjusted *p*-value (false discovery rate, FDR) below 0.05 and a minimal absolute log2 fold change (LFC) of at least 1 were considered to be differentially expressed. Normalized sequence counts were calculated accordingly with DESeq2 and log2(x+1) transformed. Transformed data (in TPM, transcripts per million) combined with previous expression data from tachyzoites, bradyzoites and enteroepithelial stages (EES [5]) are available on VEuPathDB’s ToxoDB (release 68). For all further analyses, only protein-coding genes with number of normalized counts ≥5 were considered.

The same size-factor–normalized count matrix (log₂-transformed) was re-analyzed in limma v3.46.0 (R v4.4.1) to generate intuitive stage-to-stage comparisons that complement the global DESeq2 model (full R script: Script3 Differential expression.R). A no-intercept linear model (∼ 0 + stage) was fitted, and empirical-Bayes moderation was applied to stabilize variance estimates. Contrasts were defined between each pair of consecutive developmental stages (Tachyzoite → Bradyzoite → EES1 → EES2 → EES3 → EES4 → EES5 → Unsporulated, → Sporulating → and Sporulated oocyst). Benjamini-Hochberg were employed where detection of differentially expressed genes (DEGs) was, defined by an adjusted p-value < 0.05 and an absolute log₂ fold change ≥ 2. Significant contrasts were visualized with EnhancedVolcano v1.8.0.

### Refinement of Annotation of Hypothetical Proteins

All hypothetical proteins were further annotated using OmicsBox (Version 3.2.9). Protein sequences were blasted using the blastp or blastp-fast modes against the NCBI’s nr protein database directly at NCBI using default parameters. These sequences were then mapped and annotated [11]. In parallel, InterPro domain searches [12] were performed with the same sequences. The gene ontology IDs (GOs) were finally merged. Functional annotation was expanded using eggNOG-Mapper [13].

### Variance Analysis and Clustering

To comprehensively characterize the transcriptional landscape of *T. gondii* across its life-cycle stages, an initial unsupervised clustering approach was implemented using expression data from all annotated protein-coding genes. Unless stated otherwise, analyses were performed using R (v4.4.1) and Bioconductor packages (scripts available at [Hehl Lab Toxoplasma_transcriptome_atlas](https://gitlab.uzh.ch/hehllab/toxoplasma_transcription_atlas_sporulation)). Data normalization and scaling were initially applied, retaining genes with variance <2, followed by iterative filtering to select genes with variance exceeding the mean plus one standard deviation. Dimensionality reduction was conducted on filtered genes using Uniform Manifold Approximation and Projection (UMAP) (parameters: n_neighbors = 35, min_dist = 0.04), with subsequent clustering performed via the HDBSCAN (minPts = 50). Cluster quality was quantitatively evaluated using silhouette analysis, prioritizing clusters with high silhouette widths. Mean expression profiles were computed per cluster, and biological stage identities were assigned using canonical marker gene. Visualizations, including UMAP plots, cluster expression line plots, and silhouette plots, were generated using ggplot2. Interactive UMAPs, implemented with plotly, are published at RPubs and accessible via [High-variance gene set](https://rpubs.com/Elec/HighVariance935genesTGondii) and [Full transcriptome](https://rpubs.com/Elec/FullTranscriptomicsTgondii).

### Overrepresentation Analysis (ORA)

Overrepresentation Analysis (ORA) was conducted using custom R scripts, assessing enrichment across specific life-stage contrasts: Bradyzoite vs. Tachyzoite, EES5 vs. EES2, sporulating vs. unsporulated oocyst, sporulated vs. sporulating oocyst, and unsporulated oocyst vs. gametogenesis (full script: Script4_ORA_analysis_with_cytoscape). Certain merogony comparisons (e.g., EES2 vs. EES1, EES3 vs. EES2, EES4 vs. EES3, EES5 vs. EES4) yielded minimal or no significant DEGs and were therefore excluded from ORA. Fisher’s Exact Tests were applied to determine pathway overrepresentation among DE genes, calculating log2 fold changes (the pathway’s proportion in the DE set versus its proportion in the genome) and significance thresholds (p<0.01, log2 fold change >0). Pathway analyses were performed independently for upregulated gene sets only, providing a unidirectional insight into stage-specific molecular dynamics. In addition to static bubble plots and Cytoscape exports, we generated interactive pathway–gene networks for each major stage transition.

## Results and Discussion

### Oocyst Sporulation in the Context of the Endogenous Developmental Phases

RNA-Seq analysis was performed on *T. gondii* oocysts collected from experimentally infected cats and analyzed at three defined time points during sporulation in 2% H_2_SO_4_: unsporulated (0 h), early sporulation (8 h), and fully sporulated (8 d). Microscopy confirmed the developmental stages, with no changes detected between 0 h and 8 h, but 80–81% of oocysts sporulated at 8 days (Fig. S1A). Total RNA underwent quality controls (Fig. S1B) and after mRNA enrichment, all samples were subjected to the standard RNA-Seq pipeline to produce expression data (Table S1). The reads were then mapped to the *T. gondii* reference genome of strain ME49 (ToxoDB release 68, Fig. S1C) and showed a high percentage of mapped reads to the *T. gondii* genome. Principal component analysis (PCA) of normalized gene expression revealed clear separation between fully sporulated oocysts and the earlier stages, with unsporulated and sporulating oocysts clustering closely together (Fig. S2). Differential expression analysis identified the fewest changes between unsporulated and sporulating oocysts, while >1000 genes were differentially expressed between unsporulated and fully sporulated oocysts, consistent with previous transcriptomic findings ([3], Table S2).

We combined the data newly generated transcriptomic datasetswith our previous bulk RNA-Seq data from the cat intestinal stages, tachyzoites and bradyzoites [5] to produce a comprehensive gene expression data for all the life cycle stages of *T. gondii* from the stages in the intermediate host (tachyzoites and bradyzoites), the enteroepithelial cat stages (EES1-5) and oocyst stages, considering only expressed genes with normalized reads ≥ 5 (Table S1). All genes that have not been shown to be expressed in any life cycle stage have been listed in Table S3. To refine previous annotations, we used the OmicsBox platform (Table S4). Normalization of the expression values to highest value of all life cycle stages (Table S1.4) shows clearly that many genes are stage-specific and some very specific to a single stage (Fig. 1). With this illustrative approach, we also identify a large set of genes being specific to the process of sporulation.

**Fig. 1:**
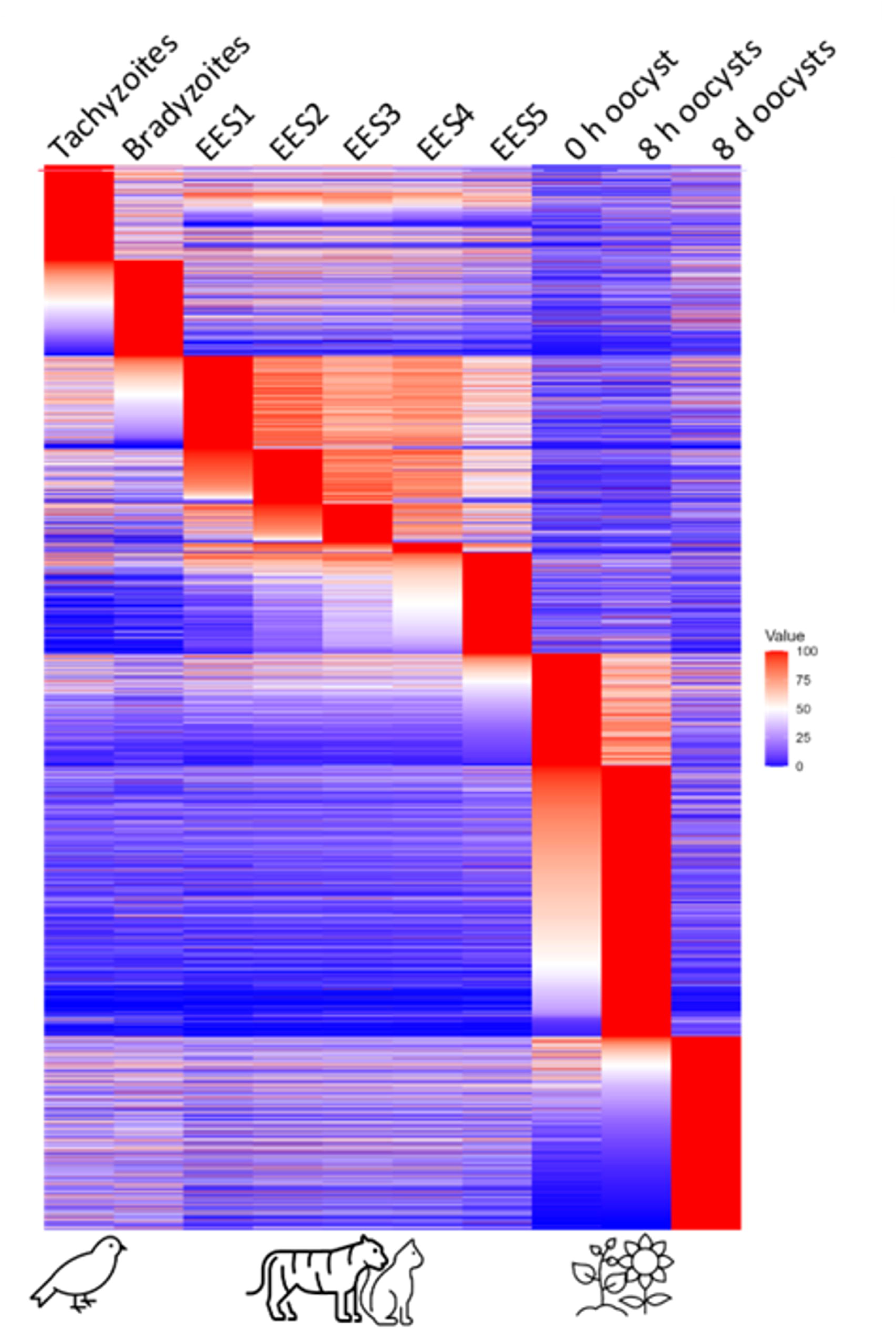
Whole protein-coding genome with gene expression values normalized to the maximum expression. Each line shows one gene. 0 h oocysts: unsporulated oocysts, 8 h oocysts: sporulating oocysts, 8 d oocysts: fully sporulated oocysts.

Building on this integrated dataset, we next employed targeted pairwise contrast analysis to precisely capture stage-specific regulatory shifts during key developmental transitions. Differential expression analysis (DEA), encompassing 7,997 protein-coding genes across ten stages (Table S5), revealed varied degrees of transcriptional remodeling (Fig. 2).

**Fig. 2.**
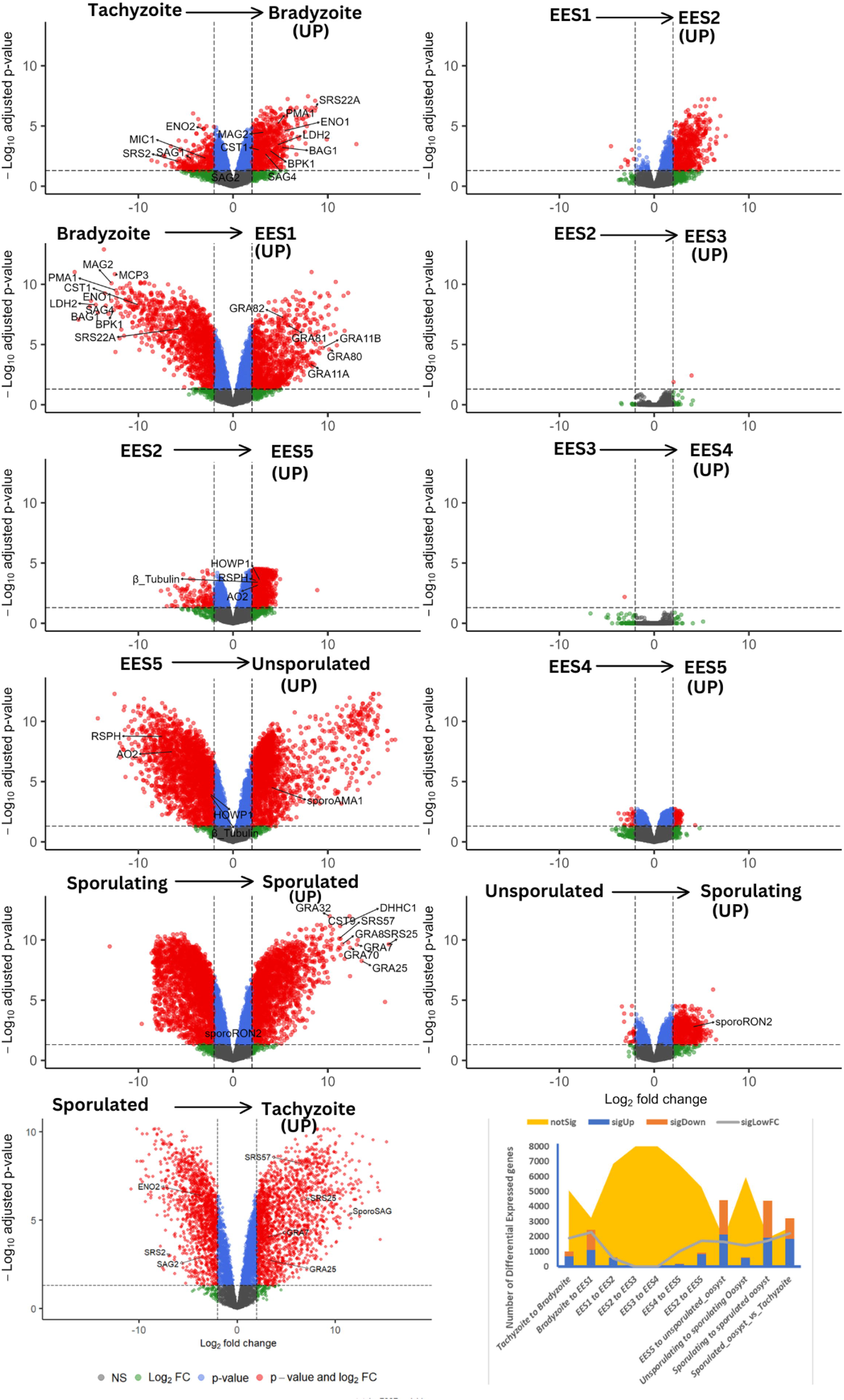
Comparative transcriptomic analyses of *T. gondii* developmental stages represented as volcano plots. The left panel (blue shaded area) highlights comparisons between closely related *T. gondii* life stages—such as EES sub-stages and transitions from unsporulated (0 h) to early sporulating (8 h) oocysts—which exhibit relatively subtle transcriptional shifts. In contrast, the right panel (orange shaded area) illustrates comparisons between more distinct developmental forms (e.g., unsporulated (8 h) vs. Sporulated (8 days) oocysts, bradyzoites vs. tachyzoites), gametocytes to oocyst, bradyzoite to EES1 revealing pronounced genetic remodeling. Each volcano plot displays differentially expressed genes, with color coding based on statistical significance and magnitude of change: Red: Genes passing both the adjusted p-value threshold (p < 0.05) and the absolute log₂ fold change criterion (≥ 2). Blue: Genes significant by p-value (p < 0.05) but not meeting the log₂ fold change threshold. Green: Genes meeting the log₂ fold change criterion (≥ 2) but not statistically significant by p-value. Gray: Genes not meeting either threshold. Known canonical markers as well as newly identified, significantly regulated genes with annotated functions are highlighted. Genes of unknown function (“hypothetical proteins”) were detected as highly significant in some comparisons but are not labeled here.

Subtle transcriptional adjustments characterized transitions among closely related enteroepithelial stages (EES) with majority of the genes remaining unchanged. Notably, minor shifts were observed between EES1 to EES2 (568 upregulated, 8 downregulated), EES2 to EES3 (2 upregulated, 0 downregulated), EES3 to EES4 (none upregulated, 1 downregulated). Moderate shifts between and EES4 to EES5 (152 upregulated, 27 downregulated), unsporulated to sporulating oocysts (595 upregulated, 24 downregulated). These incremental changes are clearly depicted by the sparsity and central clustering of significant points in the respective volcano plots.

In contrast, substantial transcriptional remodeling emerged between biologically distinct developmental stages. Pronounced gene expression changes were evident during the transition from and from EES2 to EES5 (814 upregulated, 129 downregulated), Tachyzoite to Bradyzoite (699 upregulated, 321 downregulated). Even more dramatic shifts are observed during transitions involving significant biological reprogramming: Bradyzoite to EES1 (1,109 upregulated, 1,328 downregulated), EES 5 to unsporulated oocyst (2,118 upregulated, 2,315 downregulated), sporulating to sporulated Oocyst (1,912 upregulated, 2,457 downregulated) and sporulated Oocyst to tachyzoite (1,855 upregulated, 1,352 downregulated). These prominent shifts correspond visually to the extensive dispersion and high density of significant points, reflecting both magnitude and statistical significance (Fig. 2). Comprehensive data for the upregulated or downregulated and unchanged genes can be found in supplementary (Table S5.

Taken together, this robust statistical approach combined with detailed data visualization confirms and illustrates the extent and the amplitude of the transcriptional reprogramming that accompanies major life cycle transitions in *T. gondii*, whereas more subtle incremental developmental adjustments involve few and possibly targeted gene expression adjustments. This nuanced regulatory complexity highlights promising gene candidates for further exploration of the molecular governance of stage conversion in *T. gondii*.

### Global Transcriptomic Profiling Reveals Distinct Developmental Stages in *Toxoplasma gondii*

Understanding the transcriptional changes driving *T. gondii* life cycle transitions is critical for dissecting parasite biology and pathogenicity. In *T. gondii* and other Apicomplexa, gene expression is mostly regulated on the level of transcription resulting in “just in time” expression of genes involved in cell cycle progression [14,15] and stage-differentiation. Although high-dimensional transcriptomic data capture genome-wide gene expression patterns, their complexity often obscures subtle biological signals. Traditional clustering methods tend to produce broad, heterogeneous gene sets that can mask nuanced processes. To address these challenges, we applied variance-based filtering to focus on the most useful genes, producing well-supported clusters that highlight key biological processes.

To comprehensively investigate these transcriptional dynamics, we integrated the newly generated sporulation datasets with previously published datasets covering tachyzoite, bradyzoite, and five enteroepithelial stages (EES). This integration allows, for the first time, to analyze gene expression profiles for all the life cycle stages. This is not without some caveats: although we have not seen evidence for this, we cannot exclude the theoretical possibility of a sampling bias. In addition, sampling timepoints for tissue cysts (bradyzoites) and sporulated oocysts represent local minima in the life cycle with no or only two sampling points representing transition stages, respectively. Nevertheless, our initial global transcriptomic analysis, employing an unsupervised clustering of protein-coding genes from *T. gondii* without prior filtering, aimed to capture the broad transcriptional landscape of this organism (Fig. 3a). Using UMAP reduction followed by HDBSCAN clustering, we mapped out the transcriptomic signatures across a spectrum of developmental stages. Despite the low variance threshold initially set, this approach highlighted several clusters that correspond to different biological states and transitions in the life cycle of *T. gondii.* However, a substantial number of genes could not be robustly assigned, and cluster separation was suboptimal, as indicated by low silhouette scores (Fig. S3). We then asked whether clusters could be resolved better by applying a more stringent variance-based filtering approach to the dataset. This resulted in the retention of 935 genes (Table S6) emphasizing variance as a defining feature of stage-regulated gene expression, thus providing a clearer picture of transcriptional changes throughout the life cycle (Fig. 3b). The UMAP analysis of this refined dataset containing the 935 high-variance gene set revealed well-separated clusters that were subsequently validated through silhouette analysis, showing improved internal cohesion and distinct separation of the clusters demonstrated by high silhouette scores (Fig. S4).

**Fig. 3:**
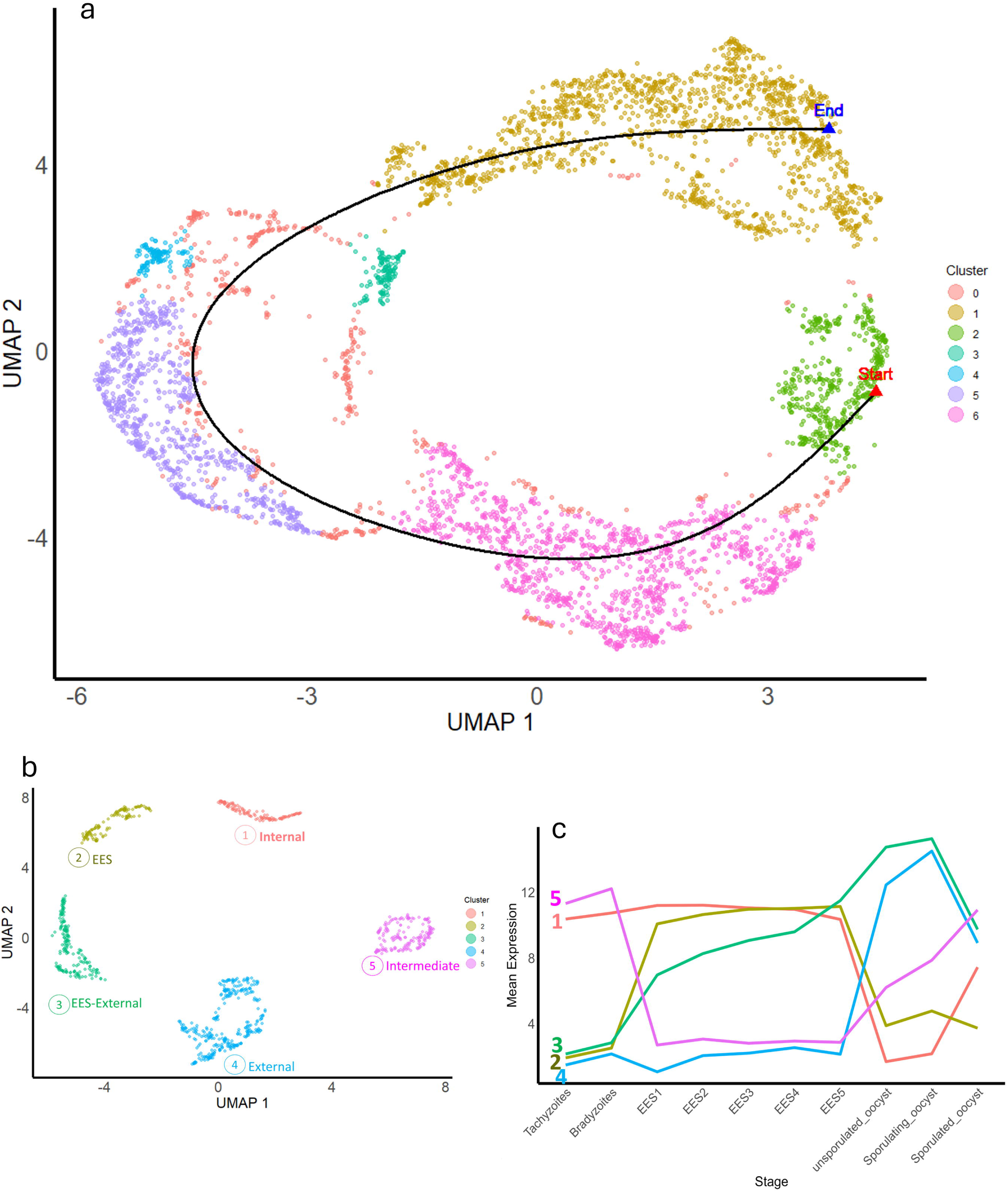
Unsupervised transcriptomic clustering across the *T. gondii* life cycle. **A** Initial UMAP embedding and HDBSCAN clustering of all protein-coding genes illustrating broad developmental transitions. **B** Improved resolution achieved by applying stringent variance-based filtering, highlighting distinct transcriptomic clusters associated with specific developmental stages: internal stages (Cluster 1), enteroepithelial stages (EES, Cluster 2), transitional stages from late EES to external (Cluster 3), external stages (Cluster 4), and intermediate-definitive host transitions (Cluster 5). **C** Mean expression profiles of these clusters across life cycle stages confirm their distinct stage-specific expression patterns.

We obtained transcript clusters that contained mixed stages. Therefore, to assign life cycle stages to the clusters, we first calculated their mean expression profiles across samples, revealing distinct patterns corresponding to specific developmental stages (Fig. 3c). For example, *Cluster 1* exhibited high average gene expression in tachyzoites, bradyzoites and all enteroepithelial stages including gametocytes. Hence, these profiles represent functions during intracellular development (both in intermediate and definitive hosts), designated as “internal”. This is in contrast to cluster 4, containing genes essentially silenced in internal stages but expressed in unsporulated, sporulating, and sporulated oocysts, designated as “external”. Interestingly, genes in cluster 2 showed a high average expression only in the enteroepithelial (EES) stage and no expression in all other stages, stages leading us to designate this cluster as “EES.” This pattern contrasts with *Cluster 5*, with expression maxima in tachyzoites, bradyzoites, in many cases beginning already during sporulation but absent in EES—thus designated as the “intermediate” (host) cluster. This cluster contains a mix of genes with expression maxima during development in the intermediate host, which represents an evolutionary more recent addition to the basic coccidial life cycle. Meanwhile, *Cluster 3* exhibited a rather distinct expression profile that highlights functions important for the core enteroepithelial and external cycle and the sporulating stages, designated as a “EES-external”.

**Table 1.** Canonical marker genes observed in this study for *Toxoplasma gondii* cluster annotation. This table summarizes canonical marker genes previously validated in the literature (transcriptomic, proteomic, or functional studies) that also show high differential expression across life stages in our dataset. These genes were used to annotate clusters from the global transcriptome, thereby validating cluster assignment to the appropriate developmental stage. (Note: additional potential markers were detected in our data but are not included here.)

| ToxoDB_ID | Product description | Gene name | Stage marker | Publication |
| --- | --- | --- | --- | --- |
| TGME49_233460 | SAG-related sequence SRS29B | SAG1 | Tachyzoites | [5], [16] |
| TGME49_268850 | enolase 2 | ENO2 | Tachyzoites | [17] |
| TGME49_291890 | microneme protein MIC1 | MIC1 | Tachyzoites | [5], [16] |
| TGME49_233480 | SAG-related sequence SRS29C | SRS2 | Tachyzoites | [5], [16] |
| TGME49_271050 | SAG-related sequence SRS34A | SAG2 | Tachyzoites | [5], [16] |
| TGME49_259020 | bradyzoite antigen | BAG1 | Bradyzoite | [18] |
| TGME49_291040 | lactate dehydrogenase | LDH2 | Bradyzoite | [19] |
| TGME49_268860 | enolase 1 | ENO1 | Bradyzoite | [20] [21] |
| TGME49_280570 | SAG-related sequence | SAG4 / SRS35A | Brdayzoite | [22] |
| TGME49_253330 | Bradyzoite pseudokinase | BPK1 | Brdayzoite | [23] |
| TGME49_264660 | SAG-related sequence SRS44 | CST1 | Brdayzoite | [20] [21] |
| TGME49_208740 | Cyst wall proteins | MCP3 | Bradyzoite | [24] |
| TGME49_252640 | P-type ATPase PMA1 | PMA1 | Bradyzoite | [25] |
| TGME49_209755 | cyst matrix protein | MAG2 | Bradyzoite | [26] |
| TGME49_238440 | SAG-related sequence | SRS22A | Bradyzoite | [27] |
| TGME49_212410 | dense granule protein | GRA11A | Merozoite | [4] [5] |
| TGME49_237800 | dense granule protein | GRA11B | Merozoite | [4] [5] |
| TGME49_273980 | dense granule protein | GRA80 | Merozoite | [28] |
| TGME49_243940 | dense granule protein | GRA81 | Merozoite | [28] |
| TGME49_277230 | dense granule protein | GRA82 | Merozoite | [28] |
| TGME49_285940 | male gamete fusion factor putative | HAP2 | EES5 | [5] |
| TGME49_233330 | radial spokehead family protein | RSPH | EES5 | [5] |
| TGME49_212240 | beta-1 tubulin, putative | $\beta$_Tubulin | EES5 | [5] |
| TGME49_500391 | copper amine oxidase, putative | AO2 | EES5 | [5] [29] |
| TGME49_316890 | hypothetical oocyst wall protein | HOWP1 | EES5 | [5] [29] |
| TGME49_315730 | apical membrane antigen | sporoAMA1 | Unsporulated | [3] [30] |
| TGME49_500398 | roptry neck protein | sporoRON2/ RON2L2 | Sporulating | [3] [30] |
| TGME49_212300 | dense granule protein | GRA32 | Sporulated | [3] |
| TGME49_250870 | palmitoyltransferase | DHHC1 | Sporulated | [31] |
| TGME49_213280 | SAG-related sequence | SRS25 | Sporulated | [32] |
| TGME49_249990 | dense granule protein | GRA70 | Sporulated | [3] |
| TGME49_203310 | dense granule protein | GRA7 | Sporulated | [3] |
| TGME49_258550 | SAG-related sequence | SporoSAG | Sporulated | [33] |
| TGME49_290700 | dense granule protein | GRA25 | Sporulated | [34] |
| TGME49_254720 | dense granule protein | GRA8 | Sporulated | [3] |
| TGME49_308020 | SAG-related sequence | SRS57 | Sporulated | [27] |
| TGME49_310790 | cyst wall protein | CST9 | Sporulated | [24] |

To further substantiate these stage assignments, we examined canonical marker genes (Table 2) associated with each developmental stage and mapped them onto the heatmap (Fig. 4). Each cluster showed unique gene expression patterns, with four clusters aligning well with known biological marker genes. For instance, *Cluster 2*—which exhibited distinct expression in all EES stages—contained GRA11A (TGME49_212410), GRA11B (TGME49_237800), GRA80 (TGME49_273980), and AO2 (TGME49_500391). These genes have been characterized as merozoite- and macro-gamete–specific markers [4,5,35]. Notably, we had previously identified GRA11A and B as merozoite markers [4]; additionally, Antunes and colleagues [35] validated the merozoites induced *in vitro* using transcriptomic analysis and identified a set of “merozoite-specific” genes—specifically the marker GRA11B [4], as well as GRA80 and GRA81 [35]. Collectively, the presence of these marker genes confirms that Cluster 2 encompasses the merozoite/pre-gamete stages. Besides merozoite marker genes, genes peaking in EES5 also belong to this cluster, e.g. TGME49_500391, encoding amine oxidase 2 (AO2), a copper amine oxidase presumably important for crosslinking of oocyst wall components [5,29]. A cluster enriched for AO2 expression would therefore align with early oocysts formation in EES5 which was shown to peak sharply in EES5. Consequently, we designated this gene *Cluster 2* the “EES” to reflect the role of these genes in *T. gondii’s* definitive host cycle.

**Fig. 4:**
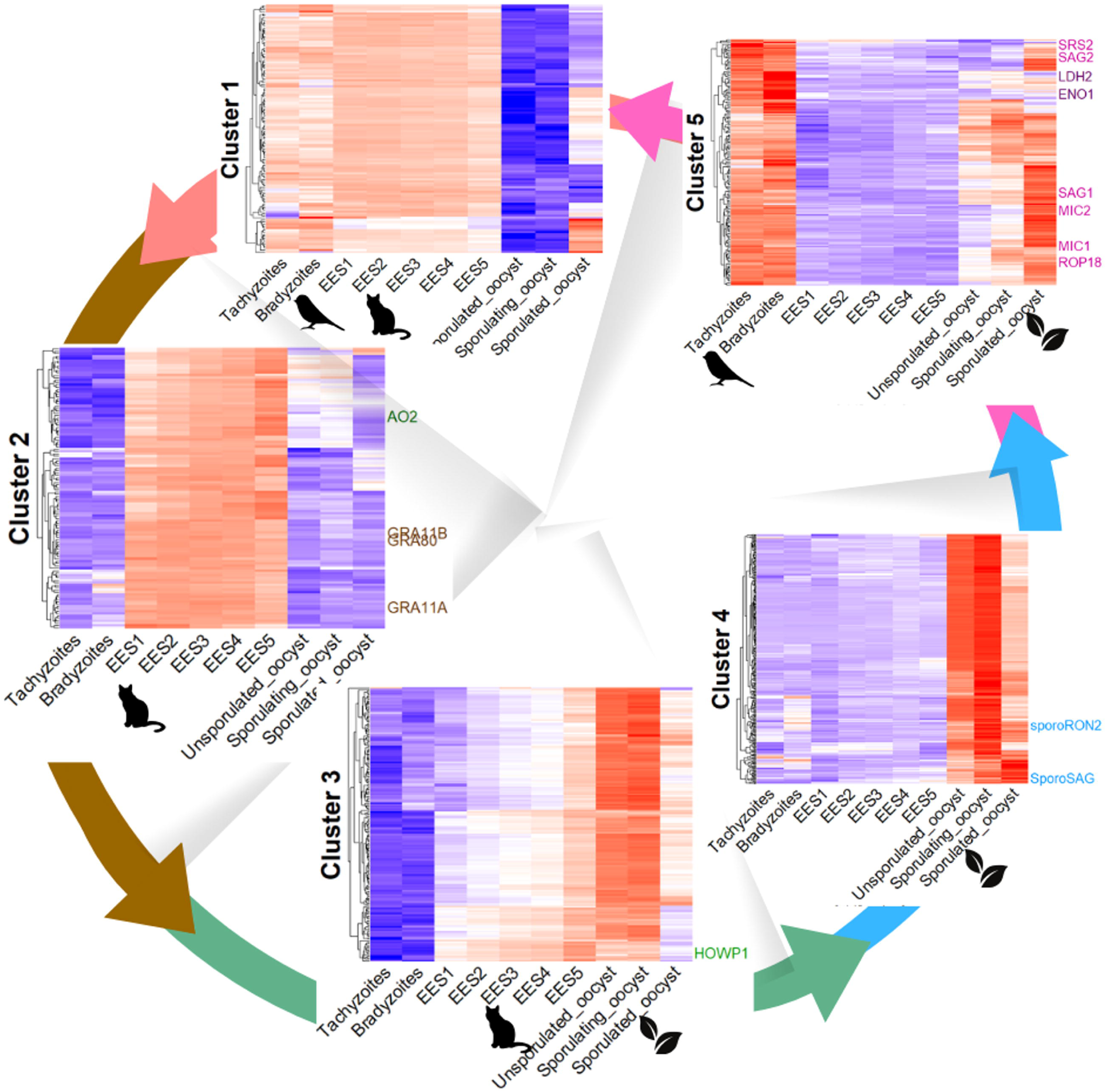
Stage-specific gene expression clusters across the *T. gondii* life cycle. Heatmaps show five major clusters based on normalized transcript abundances across tachyzoites, bradyzoites, cat enteroepithelial stages (EES1–EES5), and oocyst stages (unsporulated, sporulating, sporulated). Representative markers are highlighted: Cluster 2 (“EES stages”)—merozoite/early sexual-stage markers (GRA11A, GRA11B, GRA80, AO2); Cluster 4 (“External stages”)—sporozoite-specific genes (SporoRON2, SporoSAG); Cluster 3 (“Internal–external transition”)—gametocyte/oocyst wall– associated genes (HOWP1, β-tubulin); Cluster 5 (“Sporozoite and intermediate host stages”)—tachyzoite (SAG1, MIC1, ROP18) and bradyzoite (LDH2, ENO1) markers. Cluster name colours correspond to arrow colours in the figure to aid visual linkage. Arrows indicate developmental progression.

**Table 2:** Putative and known oocyst wall proteins with their expression maxima and relative expression values. Shaded in yellow are tyrosine-rich proteins. Data compiled from ToxoDB Release 68, Salman *et al*. [43] and Fritz *et al*. [34].i Bz: Bradyzoites, Tz: Tachyzoites, 0 h = unsporulated oocyts, 8 h = sporulating oocysts, 8 d = sporulated oocysts.

| Gene ID | Product Description | Gene Name | Tz | Bz | EES1 | EES2 | EES3 | EES4 | EES5 | 0 h | 8 h | 8 d |
| --- | --- | --- | --- | --- | --- | --- | --- | --- | --- | --- | --- | --- |
| TGME49_203500 | alanine dehydrogenase/pyridin nucleotide transhydrogenase domain-containing protein | N/A | 0 | 0 | 17 | 21 | 31 | 35 | 100 | 0 | 1 | 0 |
| TGME49_500391 | copper amine oxidase, putative | AO2 | 0 | 0 | 8 | 15 | 27 | 27 | 100 | 1 | 1 | 0 |
| TGME49_306050 | FAS1 domain-containing protein | N/A | 0 | 0 | 3 | 9 | 13 | 23 | 100 | 37 | 24 | 2 |
| TGME49_316890 | hypothetical oocyst wall protein HOWP1 | HOWP1 | 0 | 0 | 8 | 14 | 21 | 27 | 100 | 19 | 14 | 2 |
| TGME49_253150 | hypothetical protein | N/A | 0 | 0 | 13 | 18 | 33 | 29 | 100 | 0 | 0 | 1 |
| TGME49_251910 | hypothetical protein | N/A | 0 | 0 | 10 | 14 | 27 | 24 | 100 | 0 | 0 | 1 |
| TGME49_254430 | microneme protein, putative | N/A | 0 | 1 | 18 | 23 | 44 | 33 | 100 | 4 | 3 | 0 |
| TGME49_500223 | oocyst wall protein | N/A | 1 | 0 | 11 | 12 | 26 | 24 | 100 | 9 | 8 | 11 |
| TGME49_235390 | PAN domain-containing protein | N/A | 0 | 0 | 35 | 41 | 56 | 54 | 100 | 1 | 1 | 1 |
| TGME49_235200 | PAN domain-containing protein | N/A | 0 | 0 | 31 | 34 | 50 | 51 | 100 | 2 | 2 | 0 |
| TGME49_235315 | PAN domain-containing protein | N/A | 0 | 0 | 42 | 40 | 61 | 65 | 100 | 3 | 5 | 0 |
| TGME49_277572 | T. gondii family D protein | N/A | 0 | 0 | 10 | 15 | 24 | 29 | 100 | 3 | 2 | 1 |
| TGME49_294820 | type I fatty acid synthase, putative | N/A | 0 | 1 | 30 | 31 | 51 | 60 | 100 | 22 | 51 | 6 |
| TGME49_237080 | hypothetical protein | N/A | 0 | 0 | 1 | 1 | 2 | 2 | 6 | 100 | 85 | 2 |
| TGME49_258910 | hypothetical protein | N/A | 0 | 0 | 0 | 0 | 0 | 1 | 3 | 100 | 94 | 1 |
| TGME49_287510 | aromatic amino acid hydrolase AAH1 | AAH1 | 1 | 4 | 0 | 1 | 1 | 1 | 5 | 52 | 100 | 1 |
| TGME49_212740 | aromatic amino acid hydrolase AAH2 | AAH2 | 0 | 0 | 1 | 1 | 2 | 2 | 3 | 20 | 100 | 83 |
| TGME49_286250 | hypothetical protein | N/A | 0 | 0 | 0 | 0 | 0 | 0 | 3 | 72 | 100 | 1 |
| TGME49_287250 | hypothetical protein | N/A | 0 | 0 | 0 | 0 | 0 | 0 | 2 | 49 | 100 | 1 |
| TGME49_222940 | hypothetical protein | N/A | 5 | 5 | 0 | 0 | 0 | 0 | 0 | 9 | 100 | 12 |
| TGME49_320530 | hypothetical protein | N/A | 0 | 0 | 0 | 0 | 0 | 0 | 0 | 5 | 100 | 32 |
| TGME49_248730 | hypothetical protein | N/A | 0 | 0 | 0 | 0 | 0 | 0 | 0 | 5 | 100 | 4 |
| TGME49_209470 | hypothetical protein | N/A | 0 | 0 | 0 | 0 | 0 | 1 | 2 | 66 | 100 | 1 |
| TGME49_232170 | hypothetical protein | N/A | 0 | 0 | 0 | 0 | 1 | 1 | 5 | 66 | 100 | 1 |
| TGME49_269380 | hypothetical protein | N/A | 0 | 0 | 0 | 0 | 0 | 0 | 0 | 4 | 100 | 6 |
| TGME49_500127 | hypothetical protein, conserved | N/A | 0 | 0 | 0 | 0 | 1 | 1 | 4 | 53 | 100 | 1 |
| TGME49_210950 | oocyst wall protein | N/A | 0 | 0 | 0 | 0 | 0 | 0 | 0 | 4 | 100 | 4 |
| TGME49_204420 | oocyst wall protein OWP1 | OWP1 | 0 | 1 | 0 | 0 | 0 | 0 | 0 | 4 | 100 | 5 |
| TGME49_209610 | oocyst wall protein OWP2 | OWP2 | 0 | 0 | 0 | 0 | 0 | 0 | 0 | 33 | 100 | 1 |
| TGME49_268310 | oocyst wall protein OWP3 | OWP3 | 0 | 0 | 0 | 0 | 0 | 0 | 0 | 18 | 100 | 3 |
| TGME49_209180 | PAN domain-containing protein | N/A | 0 | 30 | 22 | 49 | 64 | 41 | 25 | 74 | 100 | 3 |
| TGME49_500303 | PAN domain-containing protein | N/A | 2 | 65 | 15 | 22 | 35 | 40 | 68 | 60 | 100 | 43 |
| TGME49_500061 | PAN domain-containing protein | N/A | 0 | 0 | 19 | 22 | 32 | 32 | 61 | 58 | 100 | 2 |
| TGME49_212710 | phenylalanine-4-hydroxylase | N/A | 1 | 0 | 0 | 1 | 1 | 2 | 7 | 47 | 100 | 4 |
| TGME49_271570 | T. gondii family D protein | OWP8 | 0 | 0 | 0 | 0 | 0 | 0 | 0 | 29 | 100 | 1 |
| TGME49_313000 | T. gondii family D protein | OWP9 | 0 | 0 | 0 | 0 | 0 | 0 | 0 | 4 | 100 | 12 |
| TGME49_294600 | T. gondii family D protein | OWP10 | 0 | 0 | 0 | 0 | 0 | 0 | 0 | 5 | 100 | 21 |
| TGME49_205090 | T. gondii family D protein | OWP11 | 0 | 0 | 0 | 0 | 0 | 0 | 0 | 25 | 100 | 4 |
| TGME49_272240 | T. gondii family D protein | OWP12 | 0 | 0 | 0 | 0 | 0 | 0 | 0 | 4 | 100 | 8 |
| TGME49_281590 | hypothetical protein | N/A | 0 | 0 | 0 | 0 | 0 | 0 | 0 | 2 | 9 | 100 |
| TGME49_316550 | hypothetical protein | N/A | 0 | 0 | 0 | 0 | 0 | 0 | 0 | 1 | 2 | 100 |
| TGME49_248810 | nuclear factor NF7 | N/A | 31 | 27 | 38 | 32 | 42 | 44 | 85 | 39 | 67 | 100 |
| TGME49_319890 | tyrosine-rich oocyst wall protein | TROWP | 0 | 0 | 0 | 0 | 0 | 0 | 0 | 0 | 8 | 100 |

Similarly, the distinct Cluster 4—corresponds to genes expressed maximally in extracellular parasite stages with minimal or no activity in other stages of the *T. gondii* cycle. This cluster featured well-established sporozoite markers such as SporoRON2 (TGME49_500398) and SporoSAG (TGME49_258550). Notably, SporoSAG—an SRS-family surface protein—has been directly implicated in sporozoite infectivity and is upregulated exclusively in sporozoites [3,30,33]. Systematic immunological studies in mice have shown that oral infection with sporulated oocysts elicits a distinct IgM/IgG response against recombinant SporoSAG, supporting its potential use in pinpointing the onset of sporozoite-initiated acute toxoplasmosis [33]. In parallel, transcriptomic and proteomic analyses have identified SporoRON2 together with SporoAMA1 as specialized paralogs required for sporozoite invasion; indeed, pre-incubation of sporozoites with SporoRON2 domain 3 peptide effectively blocked host cell entry [30]. In our dataset, SporoRON2 (TGME49_500398) reached peak expression at the 8 h timepoint (sporulating oocysts), and SporoSAG (TGME49_258550) peaked around 8 d (fully sporulated oocysts). Fritz et al. likewise found “SporoSAG” (SRS28) and SporoRON2 (“RON2-like2” or “RON2L2”) transcripts to be nearly undetectable at day 0 but dramatically elevated by day 4 (partially sporulated), persisting through day 10 (fully sporulated) [3]. Although the sample time points differ significantly between the two studies, both indicate that SporoRON2 and SporoSAG are effectively induced during sporulation. Taken together, these findings firmly link the high-level expression of both SporoRON2 and SporoSAG in Cluster4 to the critical roles they play during the sporozoite/oocyst stage, thereby supporting the assignment to the cluster designated “external”. Further, our dataset revealed several other genes peaking in fully sporulated oocysts, including dense granule proteins GRA32 (TGME49_212300), GRA25 (TGME49_290700), GRA70 (TGME49_249990), GRA7 (TGME49_203310), and GRA8 (TGME49_254720), as well as SRS-family members SRS25 (TGME49_213280) and SRS57 (TGME49_308020). While these proteins are not yet established as canonical sporozoite markers, their strong induction in sporulated oocysts in our data—and detection in oocyst proteomic surveys [3,27,32,34]—suggests a broader repertoire of candidate markers for the mature cyst stage.

Conversely, genes in Cluster 3 show maximum expression starting at EES5 (gametocyte-enriched) into the sporulation phase. Examples include HOWP1 (TGME49_316890), β1-tubulin (TGME49_212240) and genes coding for oocyst wall proteins (OWPs). Notably, our finding that HOWP1 peaks at EES5 aligns with our previous observations [5] where highest expression in very late *T. gondii* cat stages was observed—consistent with a macrogamete- and oocyst-specific role. Meanwhile, β-tubulin also shows marked upregulation in EES5 and hardly any expression in any other stage, mirroring its probable function in the microgamete [5] Hence, we designate Cluster 3 as an “Internal-external phase transition” cluster, reflecting how these transcripts bridge the final stages of sexual development in the cat with subsequent oocyst sporulation.

Finally, *Cluster 5* displayed gene expression profiles with maxima during intermediate host stages featuring expression of functionally or transcriptomically characterized tachyzoite markers (e.g., SAG1 (TGME49_233460), MIC4 (TGME49_208030), MIC1 (TGME49_291890), ROP18(TGME49_205250)) and bradyzoites markers (e.g. LDH2, ENO1 [5,16,19,20]). The data show increase of mRNA levels for many of these factors during sporulation. A plausible explanation is that mRNA of genes required for function in the intermediate host is already transcribed in the late sporulating stage but reaches peak expression during infection of the intracellular intermediate host. This cluster was designated “intermediate”.

Overall, this comprehensive transcriptomic analysis offers a powerful framework for unraveling and delineating distinct transcriptional signatures across *T. gondii’s* key developmental stages. By employing rigorous clustering methods and validating results with canonical stage-specific markers, we can more accurately identify the genes driving fundamental processes that define developmental stages and transitions—from tachyzoites to bradyzoites, and from enteroepithelial to oocyst stages.

### Quality Control of the RNA-Seq data

A previous transcriptomic study by Fritz *et al.* using microarray technology assessed gene expression in *T. gondii* strain M4 oocysts at various time points before and after sporulation (day 10) [3]. In the current analysis, using a lower threshold of 2-fold upregulation and a p-value < 0.05, we detected a significantly higher number of regulated genes (Fig. 5, Table S7). This is expected, given the greater read depth and sensitivity of RNA-Seq. When considering only upregulated genes in fully sporulated versus unsporulated oocysts — analogous to Fritz et al. — we detect 87% of the genes identified by microarray [3], plus more than 2700 additional genes (Fig. 5A).

**Fig. 5:**
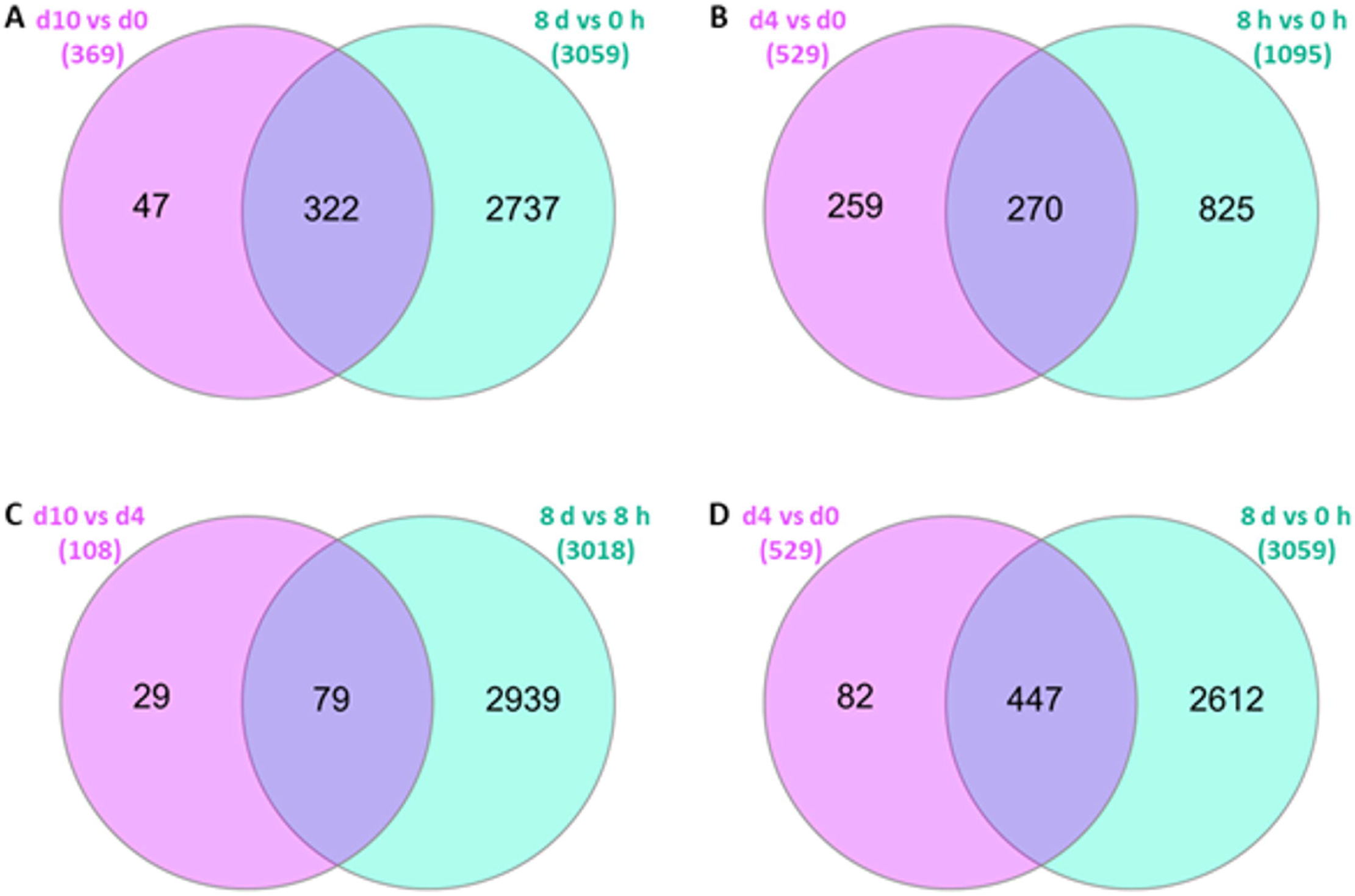
Comparison of RNA-Seq data with previous microarray data using upregulated genes in stage x versus stage y [3]. **A** Microarray d10 versus d0 and RNA-Seq sporulated (8 d) versus unsporulated (0 h). **B** Microarray d4 versus d0 and RNA-Seq sporulating (8 h) versus unsporulated (0 h). **C** Microarray d10 versus d4 and RNA-Seq sporulated (8 d) versus sporulating (8 h). **D** Microarray d4 versus d0 and RNA-Seq sporulated (8 d) versus unsporulated (0 h) For all datasets, only genes with fold changes ≥ 2 and p-values < 0.05 have been selected. All IDs are corrected for ToxoDB release 68 and only IDs existing in this release were taken into account. Venn diagrams were created using InteractiVenn [68].

In contrast, whereas the microarray study found over 500 genes upregulated at day 4 compared to unsporulated oocysts [3], our dataset reveals approximately 1100 genes upregulated in sporulating versus unsporulated oocysts (Fig. 5B), leaving almost 50% of the microarray-identified genes undetected by RNA-Seq. This discrepancy likely arises because the two different day 4 oocyst samples have been shown to contain 52% and 78% sporoblasts/sporocysts with less than half of these displaying sporozoites [3]; this suggests that these samples captured a mixture of oocysts in different phases of sporulation. This interpretation is supported by the small number of differentially regulated genes detected by microarray when comparing day 4 to day 10 oocysts, while the comparison of 8 h and 8 d RNA-Seq samples identifies >3000 upregulated genes (Fig. 5C). Furthermore, when comparing the d4 versus d0 microarray data to our sporulated (d8) versus unsporulated (0h) RNA-Seq data (Fig. 5D), we find greater overlap than when comparing to the sporulating versus unsporulated oocyst data (Fig. 5B, Table S7).

Interestingly, canonical glideosome components [36] essential for motility are upregulated in sporulated versus unsporulated oocysts, and many are also upregulated in the fully sporulated versus sporulating comparison (Table S8.1). However, in the microarray data, these genes are not consistently upregulated in the d10 sample compared with d0 and appear in the d4 versus d0 dataset again suggesting that the d4 oocysts are already highly similar to fully sporulated ones. Similarly, none of the genes encoding moving junction (MJ) components are upregulated in the microarray d10 vs. d4 data, whereas they are in our fully sporulated versus sporulating dataset (Table S8.2). Taken together, the 8-hour sampling point generated here provides a novel and informative window into the understudied process of sporulation.

In addition to the transcriptomic studies, two separate proteomics analyses of sporulated oocysts of type 2 strain M4 [34] and type II strain VEG [37] have provided an important basis on the protein composition of oocysts. Interestingly, both studies identified >100 proteins that were specific to oocysts compared to tachyzoites. We find the genes previously identified in these oocyst proteomes mainly also in combined oocyst stages or sexual stages that will include early intestinal oocysts (EES5) (Fig. S5, Table S9).

Dense granule and rhoptry proteins with expression maxima in the oocyst stages (Table S1) have not been detected in the proteomes of the oocysts [3]. This could be a result of lower sensitivity typical of proteomics compared with transcriptomics but also indicate the presence of transcripts needed for sporozoite infection in the intestine that are translationally repressed.

### Genes Coding for Oocyst Wall Proteins

*T. gondii* oocyst wall formation begins already at the haploid but mature macrogametocyte stage that contains the veil-forming-, and wall forming bodies I and II (reviewed in [38,39]). The veil is the loose outermost membrane of the maturing oocyst that is lost during shedding after the persistent oocyst walls composed of an outer and inner layer have formed underneath. During the sporulation process in the environment, two sporocysts with resistant walls containing 4 sporozoites each develop within the oocyst wall. Oocysts walls are composed mainly of proteins, acid-fast lipids, and glycoproteins and β-1,3-glucan fibrils (reviewed in [40]) with the latter being part of the inner porous layer of the oocyst wall probably providing a scaffold while the outer wall is responsible for limiting permeability [41]. The first identified cysteine-rich oocyst wall proteins (OWPs) are homologous to the *Cryptosporidium* oocyst wall proteins (COWPs) and some shown to be located at the outer wall [42]. OWPs 8-12 have been identified in later studies due to their high cysteine content rather than homology to COWPs [43]. Unexpectedly, all genes except TGME49_500223 coding for OWPs have sharp expression maxima at 8 h into sporulation (Table 2) with no or very minimal expression in EES. This shows conclusively that they are not component of the outer oocyst wall. They may, however, contribute to the oocyst wall during maturation after shedding and/or have a function in forming the sporocyst walls. Tyrosine-rich protein have been shown in *Eimeria* to be part of the inner layer of the oocyst wall [44] and in the sporocyst wall [45]; the corresponding genes display various expression maxima (Table 2) potentially. Formation of di-tyrosine bonds within or between these proteins, catalyzed by amine oxidase enzymes (e.g. AO2), confers additional stability to the oocysts and sporocysts and protect the parasite from UV radiation. The levels of AO2 mRNA peak sharply in EES5 suggesting this activity is restricted to early stages of oocyst development. The *Eimeria tenella* transcript for AO2 has been identified only in macrogametocytes whilst the protein could be detected in macrogametocytes and unsporulated oocysts [29] which is consistent with our results. At later stages during sporulation 3,4 dihydroxyphenylalanine (L-DOPA) identified in *Eimeria maxima* oocyst extracts [46] could play a role in further stabilizing the *T. gondii* oocyst wall. Interfering with genes for the two aromatic amino acid hydroxylases (AAH1: TGME49_212740, AAH2: TGME49_287510) required to convert tyrosine residues into L-DOPA and both having an expression peak in sporulating (8 h) oocysts has been shown to reduce oocyst production and sporulation [47].

Oocysts also contain Late Embryogenesis Abundant Proteins (LEAs) that are highly hydrophilic proteins and have been shown to protect oocysts against high salinity, desiccation, and freezing during long term exposure [48]. Four genes code for these proteins in *T. gondii* and are maximally expressed in sporulated oocysts (Table S1.4) consistent with their role in conferring environmental resistance.

### Canonical Sporozoite Proteins

SporoSAG is an SRS family protein detected on the surface of sporozoites [49]. Our data confirms its stage-specific expression with a sharp expression maximum in sporulated oocysts (Table S1). Sporulated oocysts express paralogues of the moving junction proteins AMA1 and RON2 called SporoAMA1 and SporoRON2 that are used in the invasion machinery [30]. Here, we show the maximum expression of the transcripts for SporoAMA1 and SporoRON2 already during sporulation (Table S1.4) potentially stockpiling for future use.

### Genome- and Life-Cycle-Wide Overrepresentation Analysis Reveals Stage-specific Metabolic Shifts During *T. gondii* Development

The addition of three transcriptomic datasets covering sporulation completes the gene expression profile of the *T. gondii* life cycle enabling a quantitative assessment of expression dynamics across all developmental stages. This comprehensive resource allows us to define the expression amplitude of individual genes and systematically track changes in cellular processes over time. To investigate how metabolic functions are modulated during stage transitions, we performed over-representation analysis (ORA) on pairwise comparisons of key life stages. ORA identifies enriched pathways, defined here as metabolic pathways statistically overrepresented among significantly upregulated genes (Fisher’s Exact Test, p < 0.01, log2 fold change > 2). An UpSet plot together with an alluvial diagram (Fig. 6) shows a quantitative representation of pathways unique to, or jointly enriched in, each of the transitions in the life cycle.

**Fig. 6:**
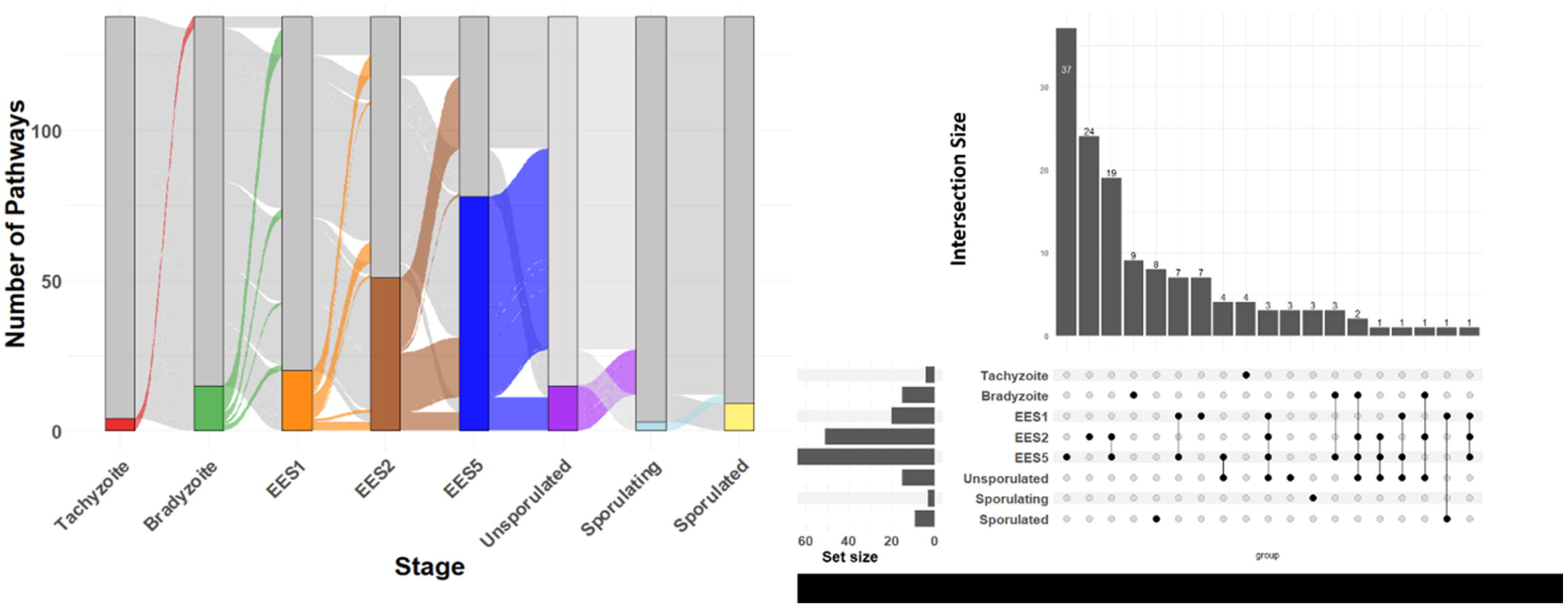
Overview of overrepresented metabolic pathways at each developmental stage of *T. gondii*. **(Left)** An **alluvial plot** shows how the total number of enriched pathways (y-axis) flows from one life-cycle stage (x-axis) to the next; each colored “stream” represents a set of pathways tracked across transitions, highlighting where they merge or diverge between stages. **(Right)** An UpSet plot quantifies unique and shared pathway enrichments across all stages. Horizontal bars (left) indicate the size (count) of enriched pathways for each individual stage (set size), while vertical bars (center) represent the intersection size for each combination of stages connected by black dots below.

Among all comparisons, transition into EES5 exhibits the highest number of significantly enriched metabolic pathways (78 at p = 0.01), with 37 unique to EES5, and the rest overlapping other stages. (Table S10). The largest overlaps occur between EES2 and EES5 (19 pathways) and between EES1 and EES5 (7 pathways). The transitions with the fewest enriched pathways are Sporulated → Tachyzoite (4) and Unsporulated → Sporulating (3)— also lack any shared pathways, indicating these sets are entirely stage-specific. The largest overlaps are concentrated in EES1 → EES2, EES2 → EES5, and EES5 → Unsporulated, while other transitions share fewer pathways (for instance, bradyzoite and EES5 overlap in only three, sporulated oocyst and EES1 in just one, and tachyzoite plus sporulating oocyst have none. These findings illustrate that while EES5 (and its neighboring transitions) show the largest array of enriched pathways, many are shared with earlier or later stages, reflecting a continuous metabolic expansion through the enteroepithelial phase. In contrast, transitions like Sporulated → Tachyzoite or Unsporulated → Sporulating feature some unique, stage-specific pathways, emphasizing the selective metabolic reprogramming that occurs at the extreme ends of the parasite’s life cycle.

We further examined each transition in turn, emphasizing how *T. gondii* reconfigures core carbon metabolism, amino-acid biosynthesis, detoxification routes, and specialised pathways to meet the distinct nutritional and physiological demands of each stage.

### Transition from Tachyzoite to Bradyzoite Fueled by Lipid and Amino Acid Catabolism, and Glycoprotein Biosynthesis

In fully established *T. gondii* bradyzoites in tissue cysts, we observe a switch to alternative energy sources—most notably lipid and branched-chain amino acid catabolism—to sustain long-term persistence [50]. In addition, *T. gondii* bradyzoites exploit xenobiotic and detoxification pathways as well as glycoprotein biosynthesis and cyst wall assembly processes. As shown in the network diagram (Fig. 7, [NDEX 1](https://www.ndexbio.org/viewer/networks/b977c071-88e0-11f0-a218-005056ae3c32)), amino acid catabolism, lipid metabolism, and xenobiotic pathways form an interconnected cluster with shared genes, whereas pathways driving glycoprotein remodeling (e.g., dolichol phosphate biosynthesis, phosphatidylglycerol degradation, tRNA splicing) appear separately regulated. Overrepresented pathways such as triacylglycerol degradation, butanoate metabolism, valine/leucine/isoleucine (VLI) catabolism, and retinoid biosynthesis support reduce replication rates while providing energy and biosynthetic precursors (largely amino acids and nitrogen) necessary for cyst maintenance. Functionally, it has been reported that bradyzoites actively scavenge host fatty acids and sequester them in lipid droplets, facilitated by enzymes like acyl-CoA:diacylglycerol acyltransferase (DGAT) [51]. Inhibition of DGAT disrupts lipid droplet formation and leads to malformed cysts, underscoring the importance of triglyceride storage for bradyzoite development [52]. These results are consistent with our data showing upregulation of lipid metabolic pathways, suggesting that bradyzoites divert resources into lipid uptake and synthesis.

**Fig. 7:**
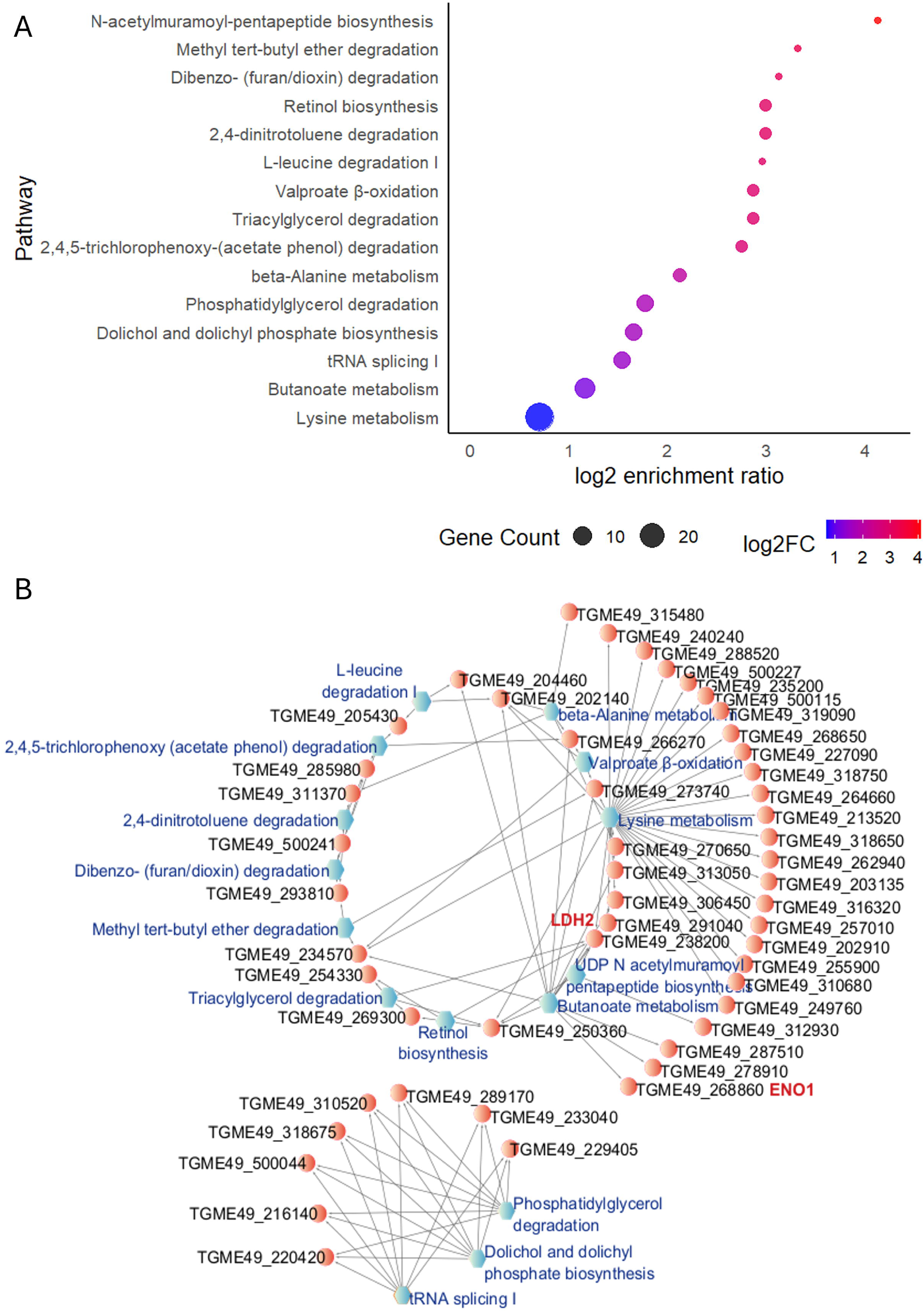
Pathway enrichment and interactive network for *T. gondii* Tachyzoite → Bradyzoite transition. (Top) Bubble plot showing the top 20 overrepresented pathways ranked by log₂ enrichment ratio. Bubble size reflects the number of genes per pathway; color scale indicates log₂ fold change. (Bottom) Cytoscape network view illustrating pathway–gene relationships, with enriched pathways (blue) connected to their associated DE genes (red labels highlight canonical markers). The interactive version of this network is available at [NDEX 1](https://www.ndexbio.org/viewer/networks/b977c071-88e0-11f0-a218-005056ae3c32).

On the other hand, the upregulation of xenobiotic-like pathways (e.g., methyl tert-butyl ether degradation, 2,4,5-trichlorophenoxy-(acetate phenol) degradation, valproate β-oxidation, 2,4-dinitrotoluene degradation) highlights an intensified focus on detoxification and antioxidant defenses, reflecting the parasite’s need to endure host immune pressures in a semi-dormant state. Transcriptome analyses have revealed that bradyzoites significantly increase expression of stress-response genes, many of which overlap with detoxification functions [53]. In one RNA-Seq study, transcripts related to heat shock, oxidative stress, and other stimuli were twice as abundant in bradyzoites compared to tachyzoites [53]. Our current dataset showing enrichment of xenobiotic metabolism pathway consistent with these observations likely reflects the upregulation of parasite genes that neutralize toxins or stress byproducts during encystation.

Additionally, ORA analysis points to repurposed processes such as UDP-N-acetylmuramoyl-pentapeptide biosynthesis—though typically linked to bacterial cell wall assembly—may contribute to cyst wall remodeling or surface glycoprotein functions. Proteomic and histochemical analyses have shown that the bradyzoite cyst wall is heavily glycosylated [54]. Accordingly, previous transcriptomic studies consistently report increased expression of genes involved in glycoprotein assembly during bradyzoite differentiation [53]. Similar trends emerged in our current transcriptomic and ORA dataset, notably identifying gene products such as tetratricopeptide repeat-containing protein ANK1 (TGME49_216140) and several cyclic nucleotide signaling enzymes including adenylate cyclase alpha 1 (TGME49_289170), 3’5’-cyclic nucleotide phosphodiesterases PDE4 (TGME49_229405), PDE5 (TGME49_220420), PDE8 (TGME49_318675), PDE12 (TGME49_310520), PDE15 (TGME49_233040), and PDE16 (TGME49_500044) associated with enriched pathways including dolichol and dolichyl phosphate biosynthesis, phosphatidylglycerol degradation, and UDP-N-acetylmuramoyl-pentapeptide biosynthesis. Although cyclic nucleotide signaling enzymes (adenylate cyclases and phosphodiesterases) are known to function in intracellular signaling rather than glycan assembly, their enrichment within these pathways might indicate previously unrecognized roles controlling glycoprotein synthesis or trafficking essential for cyst wall formation. Indeed, prior work emphasizes the critical role of dolichol-linked glycan assembly in bradyzoites, exemplified by the nucleotide-sugar transporter TgNST1 (TGME49_267380), which supplies essential sugars for glycoprotein biosynthesis required for cyst formation and long-term persistence [55]. Disrupting TgNST1 or cyst wall glycoproteins, such as proteophosphoglycan (TgPPG1) and CST1 (a glycoprotein containing *N*-acetylgalactosamine (GalNAc) [56], compromises cyst structure and bradyzoite viability [55,57,58]. Although both are essential for cyst wall formation, TgPPG1 mRNA [57] is expressed at all life cycle stages whilst CST1 expression is restricted to bradyzoites [58], reflecting coordinated glycoprotein production during encystation [57]. Collectively, these findings illustrate a metabolic pivot from energy-intensive proliferation toward catabolically flexible, and conservation-oriented mechanisms, ensuring *T. gondii*’s long-term survival in cysts within host tissues.

### Metabolic Remodeling from Bradyzoites to EES1: Upregulated Lipid, Core Carbon, Sulfur, and Aromatic Amino Acid Pathways in *T. gondii*

The tissue cyst can be viewed as a local minimum in the life cycle which requires activation to emerge from its metabolic quiescent state. Upon relief of immunological pressure bradyzoites differentiate to tachyzoites in all cells of all hosts except feline enterocytes. Only in these cells a gene expression program is launched resulting in development of the very early enteroepithelial stage (EES1). Development of rapidly dividing merozoites triggers major metabolic shifts involving lipid metabolism, central carbon metabolism (TCA and glyoxylate cycles), sulfur and seleno-amino acid metabolism, and aromatic amino acid biosynthesis (Fig. 8, [NDEX 2](https://www.ndexbio.org/viewer/networks/b0998dd3-8a2d-11f0-a218-005056ae3c32)). Specifically, our analyses revealed pronounced upregulation of the TCA cycle, C5-branched dibasic acid metabolism, and L-glucose degradation, indicative of increased requirements for biosynthetic intermediates and flexible carbon utilization as bradyzoites reactivate into enteric merozoites. This metabolic transition aligns with previous transcriptomic observations showing upregulation of glycolytic and TCA cycle genes in merozoites [59]. Similar metabolic patterns have been documented in other coccidian parasites, such as *Eimeria* which exhibit elevated expression of glycolytic transcripts in merozoites and gametes relative to quiescent sporozoite stages [60]. At the same time, our findings demonstrate activation of fatty acid biosynthesis (type II) and glyoxylate cycle/fatty acid degradation superpathways. Indeed, prior studies have also reported increased expression of genes linked to fatty acid biosynthesis pathways, largely inactive during encystation but activated upon parasite reactivation; conversely, fatty acid β-oxidation, active in cyst stages, is suppressed following reactivation [59]. The simultaneous activation of fatty acid biosynthesis and glyoxylate pathways emphasizes increased demands for membrane biogenesis and energy production at this stage.

**Fig. 8.**
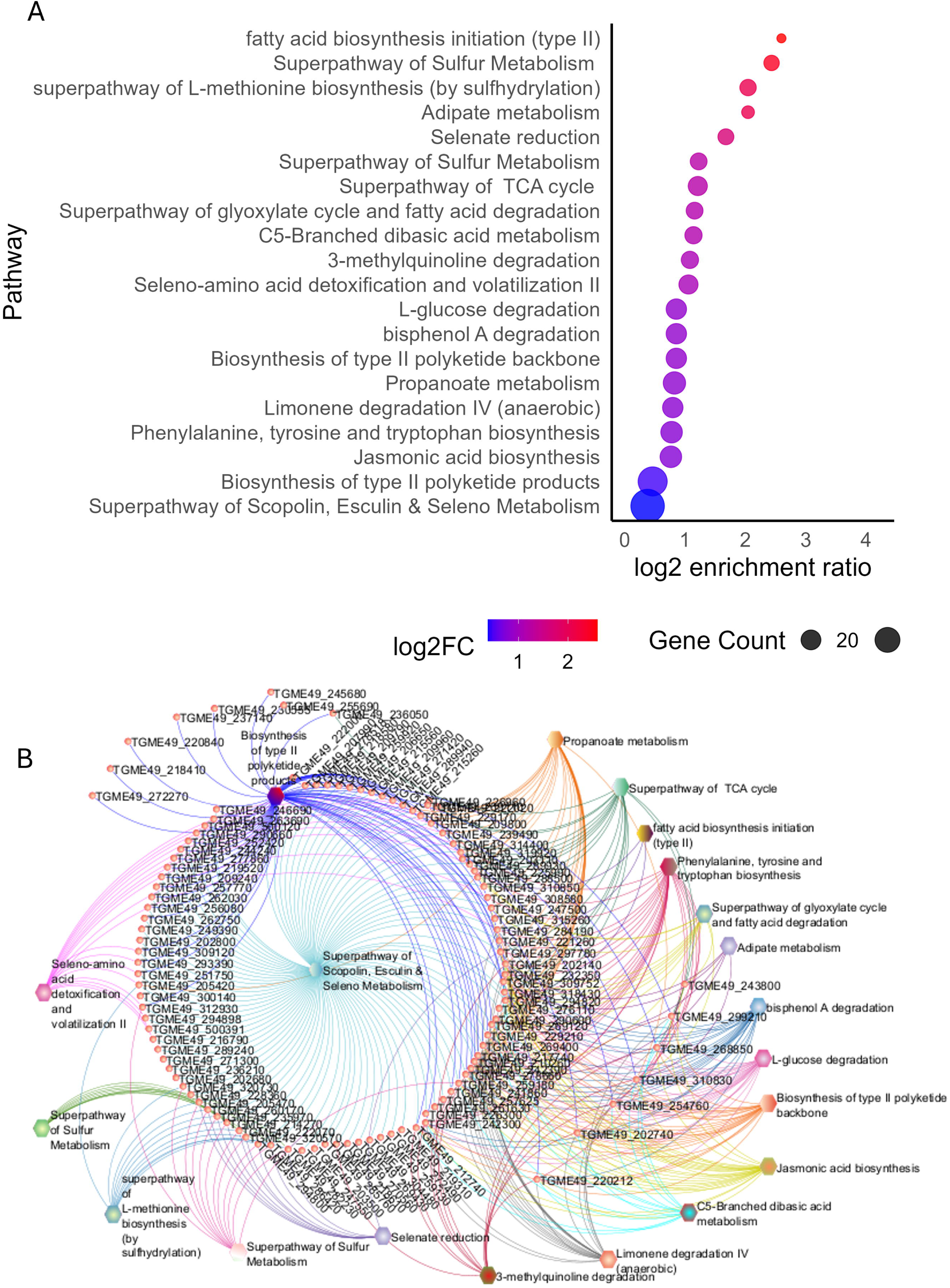
Pathway enrichment and interactive network for the *T. gondii* Bradyzoite → EES1 transition. (Top) Bubble plot showing the top enriched pathways ranked by log₂ enrichment ratio. Bubble size reflects the number of genes per pathway; color scale indicates log₂ fold change. (Bottom) Cytoscape network view illustrating pathway–gene relationships, with enriched pathways (colored nodes) connected to their associated DE genes. Prominent pathways include sulfur metabolism, fatty acid biosynthesis, polyketide biosynthesis, and selenate reduction, highlighting major metabolic rewiring during the onset of merogony. The interactive version of this network is available at [NDEX 2](https://www.ndexbio.org/viewer/networks/b0998dd3-8a2d-11f0-a218-005056ae3c32).

Furthermore, our data highlight significant induction of sulfur metabolism, including selenate reduction and L-methionine biosynthesis pathways. Although *T. gondii* possesses a complete reverse trans-sulphuration pathway for cysteine biosynthesis [61], stage-specific upregulation of sulfur metabolism has not previously been reported. Similarly, the genomic capacity of *T. gondii* for selenium detoxification via cysteine desulfurase/selenocysteine lyase has not been previously associated with developmental transitions, making the observed upregulation of seleno-amino acid metabolism particularly noteworthy. Additionally, enrichment of phenylalanine, tyrosine, and tryptophan biosynthesis pathways was observed. Aromatic amino acid hydroxylases (AAH1/2), known for producing L-DOPA, are elevated in bradyzoite cyst stages, facilitating initiation of merogony in cat enterocytes [62]. Our findings extend this observation to the EES1 stage, emphasizing a potential broader role for aromatic amino acids in proteome remodeling during early enteroepithelial infection.

Notably, enrichment of type II polyketide and jasmonic acid biosynthesis pathways was also detected. Given that *T. gondii* harbors large polyketide synthase (PKS) genes reminiscent of bacterial PKSs capable of synthesizing secondary metabolites [62], these findings suggest previously unexplored roles for specialized metabolites in supporting rapid parasite growth and internal environment adaptation during the EES1 stage. Finally, network analysis (Fig. 8) highlights the extensive interconnectivity among these enriched metabolic pathways, illustrating how individual metabolic modules synergistically sustain the complex demands of the parasite during the critical establishment of enteroepithelial stages. Overall, the coordinated enrichment and novel interconnectivity of these metabolic pathways revealed by our study present an integrative and unprecedented perspective, extending significantly beyond the scope of previous research.

### Coordinated Metabolic Remodeling of *Toxoplasma gondii* from EES1 to EES2 and Beyond: Sulfur Assimilation, Carbon Metabolism, and Amino Acid Biosynthesis Extending into EES5

As *T. gondii* transitions from EES1 to EES2, it significantly expands its metabolic repertoire, particularly in sulfur assimilation, carbon metabolism, and amino acid biosynthesis, as demonstrated by pathway enrichment and network analyses of our dataset (Fig. 9, [NDEX 3](https://www.ndexbio.org/viewer/networks/a8c7164d-1611-11f0-9806-005056ae3c32)). Our analyses revealed notable enrichment and interconnectivity among sulfur-related pathways and amino acid biosynthetic processes, including the biosynthesis of L-threonine, mimosine, taurine, L-cysteine, the interconversion of homocysteine and cysteine, and hydrogen sulfide production. Key genes associated with methionine metabolism—such as TGME49_259180 (pyridoxal-phosphate-dependent superfamily protein), TGME49_278910 (homoserine O-acetyltransferase, putative), TGME49_320730 (O-acetylserine (thiol) lyase 2, putative), and TGME49_325200 (cystathionine beta-synthase)— exhibited marked upregulation, underscoring a robust enhancement of sulfur utilization coincident with increased amino acid production during early enteric development. Our previous transcriptomic comparing *T. gondii* merozoites to tachyzoites [4] suggested upregulation of the methionine biosynthesis pathway, primarily evidenced by increased expression of homoserine O-acetyltransferase. However, comprehensive activation of broader sulfur assimilation processes, such as L-cysteine biosynthesis and hydrogen sulfide synthesis, had not been previously documented. Thus, our findings represent the first direct evidence of extensive sulfur pathway activation during the early EES transition.

**Fig. 9:**
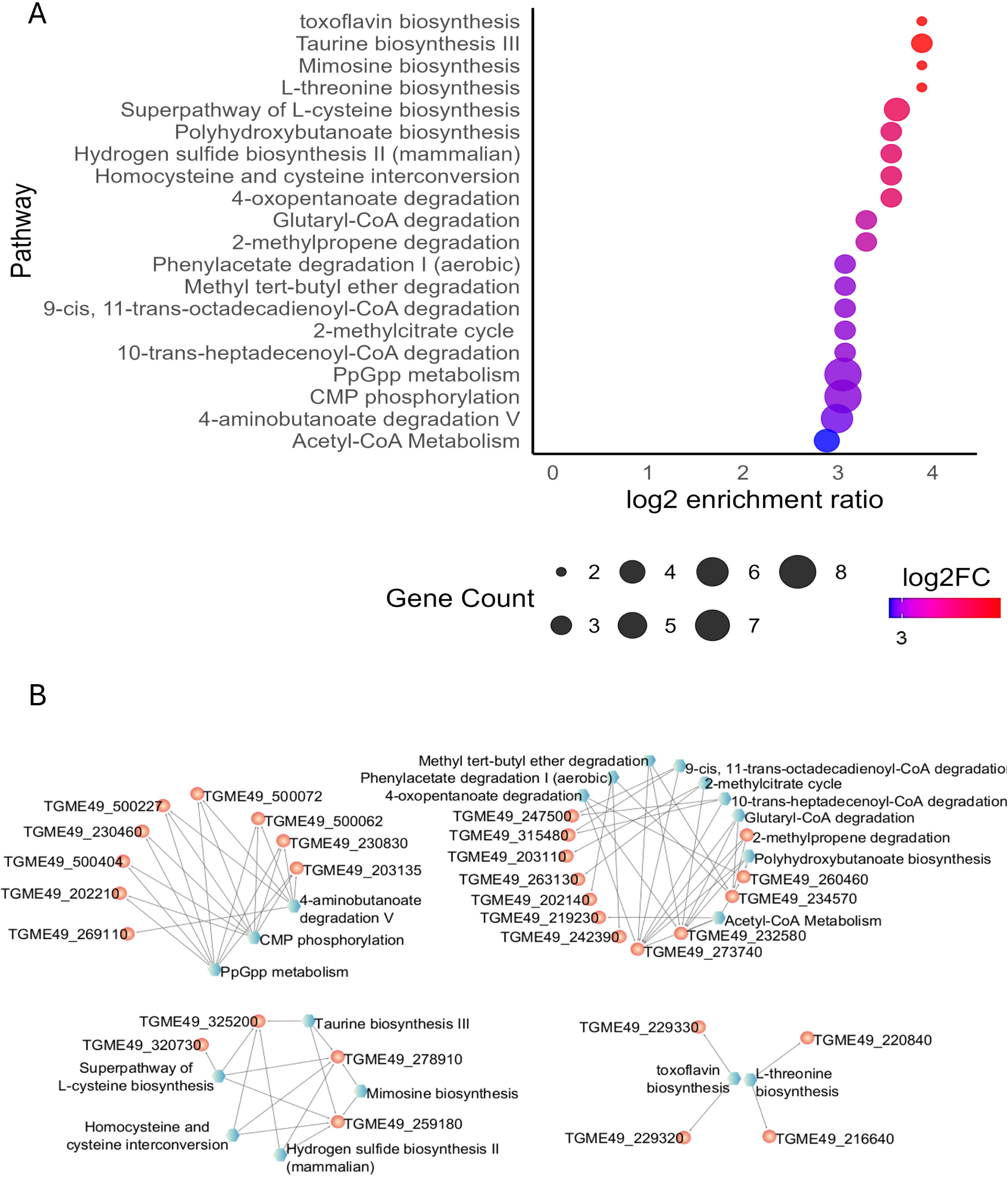
Pathway enrichment and interactive network for the *T. gondii* EES1 → EES2 transition. (Top) Bubble plot showing the top enriched pathways ranked by log₂ enrichment ratio. Bubble size reflects the number of genes per pathway; color scale indicates log₂ fold change. (Bottom) Cytoscape network view illustrating pathway–gene relationships, with enriched pathways (blue labels) connected to their associated DE genes. Notable pathways include L-cysteine biosynthesis, taurine biosynthesis, hydrogen sulfide biosynthesis, and polyhydroxybutanoate biosynthesis, indicating metabolic rewiring during early enteric development. The interactive version of this network is available at [NDEX 3](https://www.ndexbio.org/viewer/networks/a8c7164d-1611-11f0-9806-005056ae3c32).

We also detected a second network cluster indicating increased carbon utilization, lipid catabolism, and energy storage capacity—evident in pathways such as fatty-acyl-CoA degradation, 4-oxopentanoate degradation, 2-methylpropene degradation, polyhydroxybutanoate biosynthesis, and acetyl-CoA metabolism. Prominent enzymes driving these metabolic shifts included TGME49_232580 (oxidoreductase), TGME49_260460 (acyl-CoA synthetase (ACS6)), and TGME49_273740 (acetyl-CoA acyltransferase B). These findings illustrate a shift towards enhanced catabolic versatility and carbon storage in EES2. Previous RNA-Seq analyses of *T. gondii* merozoites showed moderate increases in mRNA levels of glycolytic and TCA cycle genes compared to tachyzoites corroborating our observation of intensified carbon flux in the early enteric phase [47]. However, the specific upregulation of fatty-acyl-CoA degradation and polyhydroxybutanoate biosynthesis was not documented in *T. gondii* before, although some analogous pathways were documented in *Eimeria* gametocyte stages [60]. Our findings thus expand the known repertoire of catabolic and storage processes leveraged by *T. gondii* during the enteric cycle.

In parallel, we noted a distinct cluster encompassing regulatory and specialized pathways—such as ppGpp metabolism, CMP phosphorylation, and 4-aminobutanoate degradation V. PpGpp metabolism, refers to the alarmone guanosine tetra/pentaphosphate, a signaling nucleotide in bacteria and plant chloroplasts [63]. Apicomplexan parasites harbor the non-photosynthetic plastid termed apicoplast and encode RelA/SpoT homologs that could produce (p)ppGpp, regulating metabolic gene expression in response to stress or developmental cues. The “stringent response” is well characterized in bacteria, where (p)ppGpp accumulation under nutrient stress leads to widespread metabolic reprogramming [63]. However, the involvement of an alarmone-like (p)ppGpp signaling pathway has yet to be demonstrated in *T. gondii* or other apicomplexan parasites. Consequently, our identification of putative ppGpp-related enzymes in the EES2 transcriptome suggests a hitherto uncharacterized regulatory mechanism, potentially paralleling the canonical “stringent response” found in prokaryotes.

As the parasite advances further to EES5, we observed expansions in these same metabolic pathways, now including valproate β-oxidation, ketone body metabolism, ethanol degradation, L-glutamate degradation, and L-ornithine biosynthesis. This broader metabolic toolkit underscores *T. gondii*’s capacity to exploit diversified carbon sources, facilitate nitrogen assimilation, and sustain sulfur utilization through the later stages of the enteric cycle. Previous studies in *Eimeria*, identified increased lipid catabolism and partial reliance on ketone-body-dependent pathways during sexual development [29], yet until now these processes have not been directly investigated in *T. gondii* EES5. Our data reveals similarities between *T. gondii* and *Eimeria* metabolism during gamete development but also show evidence for additional metabolic routes—such as valproate β-oxidation and ethanol degradation. Altogether, these metabolic expansions may serve the heightened demands of gametogenesis, structural remodeling, and rapid replication characteristic of the EES5 stages (Table S10).

In summary, our transcriptome and network analyses reveal broader, interconnected metabolic rewiring in *T. gondii* EES stages, enhancing utilization of diverse substrates through increased sulfur assimilation, amino acid biosynthesis, carbon degradation, and regulatory pathways. These metabolic shifts support the parasite’s escalating biosynthetic and energetic demands from early merogony (EES1–EES2) through advanced gametogenesis (EES5).

### Fatty acid, Ketone Body Metabolism and Expanded Detoxification: Metabolic Reprogramming of *Toxoplasma gondii* from EES5 to Unsporulated Oocysts

The transition from EES5, harvested in the cat intestine to the unsporulated oocyst isolated from fresh cat feces, key metabolic themes intensify, particularly emphasizing alternative energy production and detoxification pathways (Fig. 10 [NDEX 4](https://www.ndexbio.org/viewer/networks/92d45ead-8991-11f0-a218-005056ae3c32)). Among the top enriched pathways based on enrichment scores (log2 fold change) and significance are ketone body metabolism, methyl tert-butyl ether degradation, and metabolism of xenobiotics by cytochrome P450, underscoring the parasite’s specialized metabolic adaptation at this developmental stage. Previous proteomic studies in *T. gondii* oocysts noted elevated fatty acid β-oxidation and branched-chain amino acid degradation (Pizzi et al., 2013), and functional characterization demonstrated a cytochrome P450 enzyme’s detoxification role (Zhang et al., 2018). However, our transcriptomic analysis uniquely identifies ketone body metabolism genes as exclusively detected in unsporulated oocysts, revealing a novel metabolic adaptation possibly involved in acetyl-CoA buffering and energy mobilization during environmental transition.

**Fig. 10:**
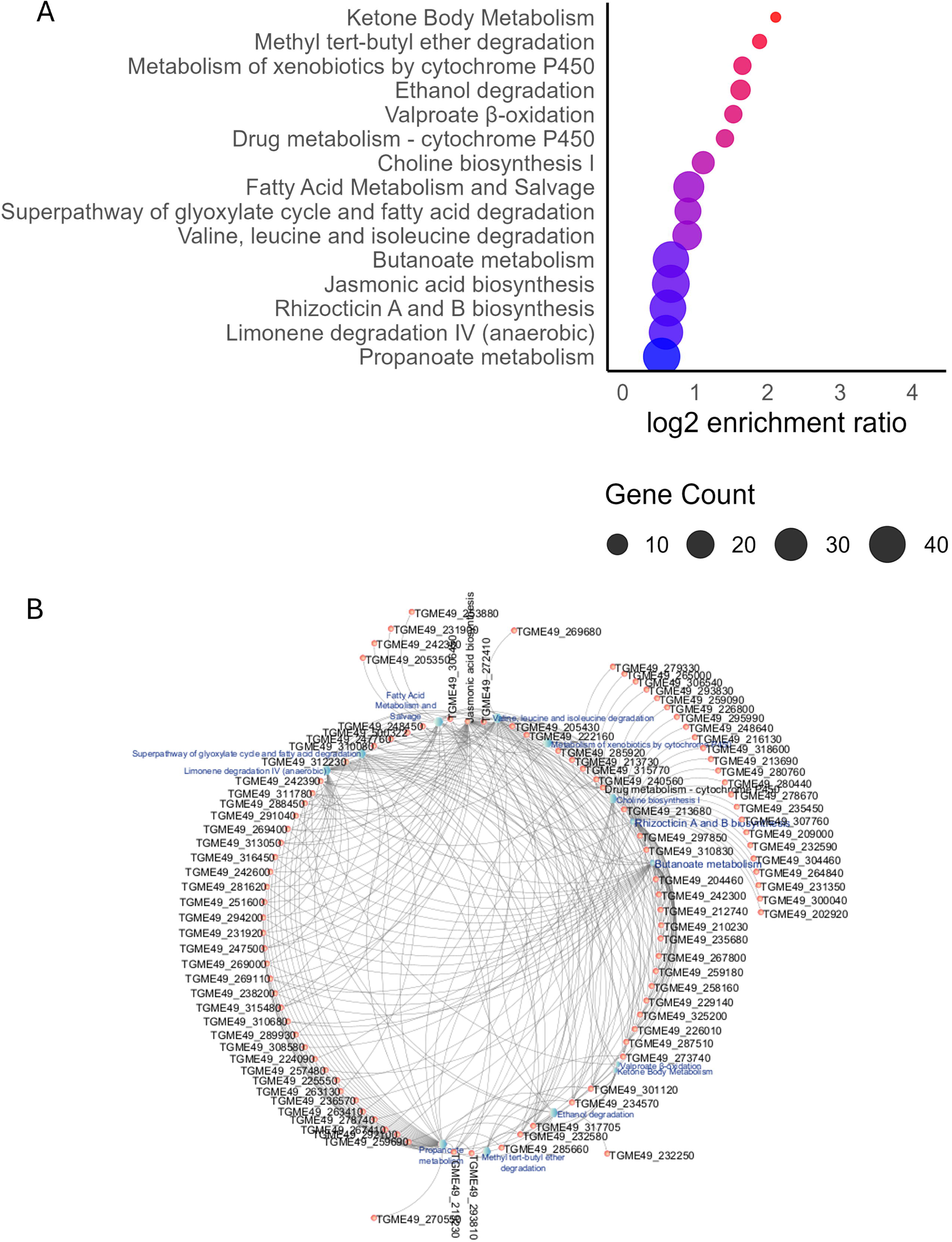
Pathway enrichment and interactive network for the *T. gondii* EES5-Unsporulated transition. (Top) Bubble plot showing the top enriched pathways ranked by log₂ enrichment ratio. Bubble size reflects the number of genes per pathway; color scale indicates log₂ fold change. (Bottom) Cytoscape network view illustrating pathway–gene relationships, with enriched pathways (blue labels) connected to their associated DE genes. Prominent pathways include ketone body metabolism, cytochrome P450–mediated xenobiotic metabolism, and fatty acid degradation, consistent with enhanced metabolic plasticity during sporulation onset. The interactive version of this network is available at [NDEX 4](https://www.ndexbio.org/viewer/networks/92d45ead-8991-11f0-a218-005056ae3c32).

Additional significantly enriched pathways include ethanol degradation, valproate β-oxidation, drug metabolism via cytochrome P450, and choline biosynthesis, reinforcing the parasite’s robust detoxification capabilities and nutrient reallocation strategies. While previous work has documented cytochrome P450-related detoxification, specific enrichment in xenobiotic degradation pathways such as methyl tert-butyl ether and valproate β-oxidation has not been reported before, marking these as new insights into the parasite’s expanded metabolic repertoire. Furthermore, our findings support prior evidence from studies in related coccidia like *Eimeria,* which show a similar metabolic shift emphasizing lipid and amino acid substrates to fuel sporulation and environmental persistence (Walker et al., 2015; Fritz et al., 2012).

The network diagram (Fig. 10, [NDEX 4](https://www.ndexbio.org/viewer/networks/92d45ead-8991-11f0-a218-005056ae3c32)) reveals substantial interconnectivity, with many shared genes bridging lipid metabolism, branched-chain amino acid catabolism, and detoxification pathways. Notably, ketone body metabolism emerges as uniquely specialized: the four annotated genes—acetyl-CoA acetyltransferase (TGME49_204460), putative acyl-CoA synthetase ACS6 (TGME49_232580), putative acetyl-CoA acyltransferase B (TGME49_273740), and putative hydroxymethylglutaryl-CoA lyase (TGME49_301120)—were exclusively expressed in the unsporulated oocyst. This specificity strongly indicates a previously unreported pivotal role for these genes in nutrient storage, detoxification, and metabolic reallocation strategies during the environmental adaptation of the parasite. The identification of enriched unconventional pathways such as anaerobic limonene degradation, rhizocticin biosynthesis, jasmonic acid biosynthesis, and propanoate metabolism further highlights *T. gondii*’s metabolic plasticity, indicating enzymes with broad substrate specificity and uncovering previously unrecognised biochemical capabilities at this critical life stage.

### Specialized Secondary Metabolism and Environmental Resilience in Fully Sporulated *Toxoplasma gondii* Oocysts: Terpenoid, Menaquinone, and Redox Pathway Enrichment

As sporulation is completed, the parasite attains a second local minimum with no cell division and metabolic reprogramming underscoring a transition from active developmental remodeling toward enhanced environmental resilience. Specialized secondary metabolic pathways are enriched in fully sporulated oocysts, notably including those involved in terpenoid and menaquinone biosynthesis, along with mechanisms linked to redox homeostasis such as selenate reduction (Fig. 11, [NDEX 5](https://www.ndexbio.org/viewer/networks/f1b8ba87-8980-11f0-a218-005056ae3c32)). These findings suggest an adaptive transition favoring survival outside the host environment.

**Fig. 11:**
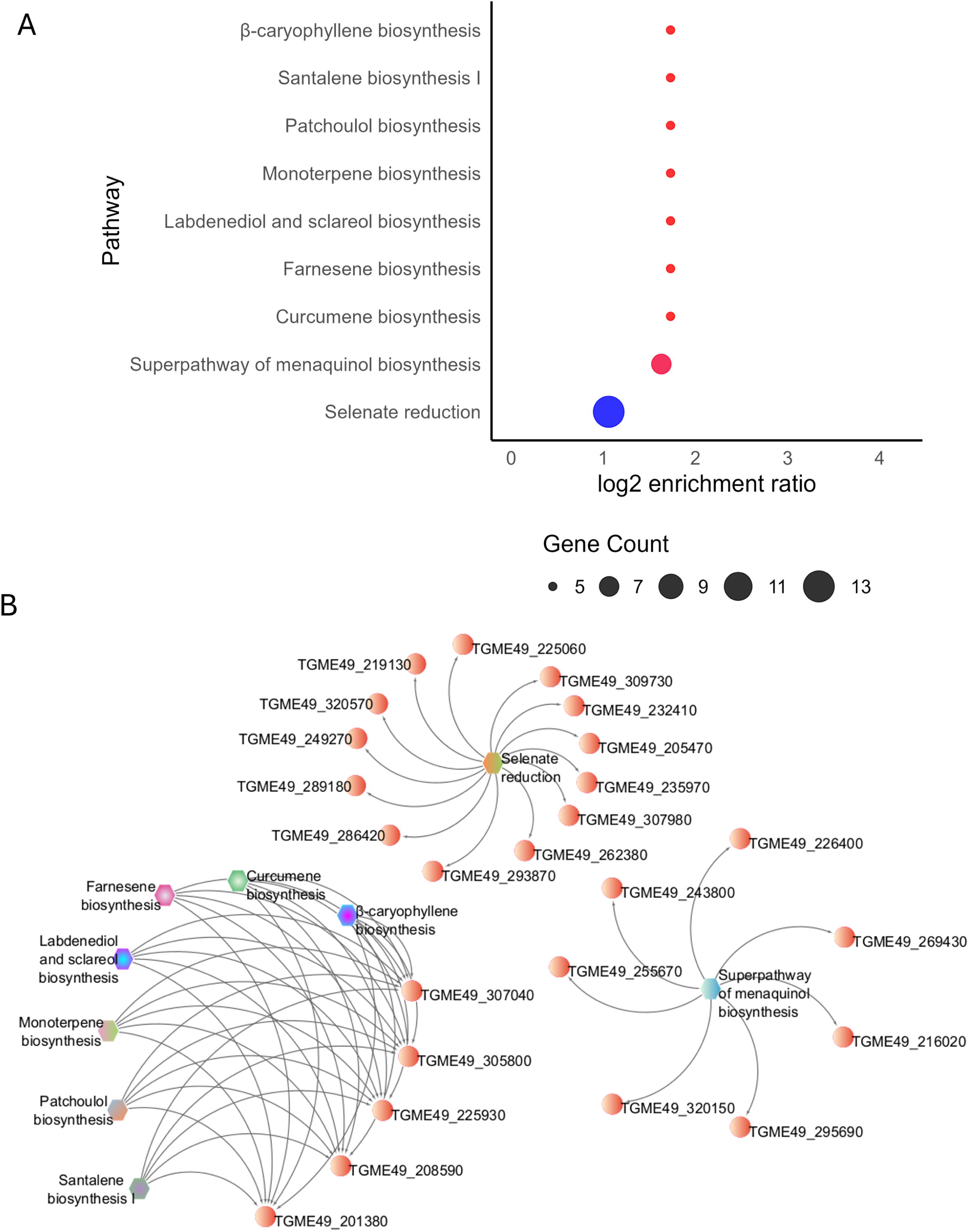
Pathway enrichment and interactive network for the *T. gondii* Sporulating → Sporulated transition. (Top) Bubble plot showing the top enriched pathways ranked by log₂ enrichment ratio. Bubble size reflects the number of genes per pathway; color scale indicates log₂ fold change. (Bottom) Cytoscape network view illustrating pathway–gene relationships, with enriched pathways (colored nodes) connected to their associated DE genes. Highlighted pathways include terpenoid biosynthesis, menaquinone biosynthesis, and selenate reduction, reflecting specialized secondary metabolism and redox adaptations in fully sporulated oocysts. The interactive version of this network is available at [NDEX 5](https://www.ndexbio.org/viewer/networks/f1b8ba87-8980-11f0-a218-005056ae3c32).

Specifically, top-ranking pathways in our analysis include the superpathway of menaquinol biosynthesis and selenate reduction. Such enrichment strongly points toward an enhanced capacity for specialized electron transport, antioxidant defense, and detoxification mechanisms. Similarly, terpenoid biosynthetic pathways (β-caryophyllene, santalene, patchoulol, monoterpene farnesene, curcumene, labdenediol and sclareol biosynthesis, are significantly represented, indicating broader adaptations aimed at producing protective secondary metabolites. The abundance of terpene-associated pathways (e.g., santalene, patchoulol, labdenediol, and sclareol biosynthesis) likely reflects specialized or ancestral plastid-related functions that reinforce the environmental stability and robustness of the sporulated oocysts. Indeed, functional studies have demonstrated the metabolic importance of the apicoplast organelle and related plastid pathways in *T gondii*, particularly concerning the parasite’s survival and persistence [47,64,65]. Additionally, the transcriptomic study by Fritz *et al.* [3] identified unique metabolic features in *T. gondii* oocysts, suggesting the involvement of plastid pathways in the parasite’s environmental resilience. Collectively, these studies provide evidence that the apicoplast and its associated metabolic functions play an important role to the survival and persistence of *T. gondii* oocysts in external environments.

Interestingly, our enrichment data (Table S10) indicates that these specialized pathways are each represented by only a small subset of genes in the *T. gondii* genome. Remarkably, nearly all of these genes rank among the most highly upregulated in fully sporulated oocysts. For example, the biosynthetic pathways for curcumene, farnesene, labdenediol/sclareol, monoterpene, patchoulol, and santalene each contain six annotated genes. Five out of six are consistently upregulated in fully sporulated oocysts including: 6-pyruvoyl tetrahydrobiopterin synthase (TGME49_201380), chorismate synthase, putative (TGME49_208590), a shikimate dehydrogenase substrate binding domain-containing protein (TGME49_225930), a vacuolar ATP synthase subunit 54 kDa, putative (TGME49_305800), and triose-phosphate isomerase TPI-I (TGME49_307040). Such precise activation underscores the highly specialized nature of these metabolic pathways, indicating that these routes are not broadly constitutive but rather dormancy- or survival-specific genetic circuits selectively activated during sporulation. This robust enrichment highlights their potential roles in stress tolerance, environmental protection, and maintaining infectivity once the oocyst is released into the external environment.

Previous transcriptomic, proteomic, and functional analyses of *T. gondii* oocysts have primarily emphasized broad-scale transcriptional and functional changes during sporulation, identifying general stress-response pathways, antioxidant enzymes, and oocyst-wall protein genes [3,34,42,47,48]. For example, Fritz and colleagues [3] characterized global patterns of gene expression changes but did not explicitly dissect the roles of specific secondary metabolic pathways or plastid-related metabolic routes during sporulation. Similarly, proteomic studies reported broad enrichments in metabolic and translation-related processes [47]. Building on this, Arranz-Solís provided more targeted insights by functionally characterizing the critical role LEA proteins play in stress tolerance [48]. Yet, the functional implications of secondary metabolic enzymes, terpenoid synthesis, or specialized redox pathways have remained largely unexplored. Notably, comparative analyses of related apicomplexans such as *Eimeria* and *Cryptosporidium* have shown parallels in stress-response gene activation and oocyst-wall formation, but highlight distinct metabolic capabilities likely driven by lineage-specific life-cycle adaptations, such as the absence of apicoplast-related terpenoid biosynthesis in *Cryptosporidium* [66,67].

Our current findings explicitly address these gaps by identifying transcriptomic enrichment of specialized secondary metabolic pathways potentially underpinning the environmental durability of fully sporulated *T. gondii* oocysts. By highlighting these selectively activated metabolic routes, our transcriptome-based ORA analysis significantly expands the understanding of adaptive metabolic strategies *T. gondii* may employ to ensure environmental survival and future infectivity. The observed enrichment of terpenoid backbone biosynthesis, menaquinone/ubiquinone metabolism, and selenate reduction provides clear examples of metabolic priming previously unrecognized at the transcriptional level in mature oocysts, positioning our findings as novel insights complementing existing transcriptomic and proteomic literature [3,47,48]. Future studies may focus on experimental validation of these pathways, exploring the regulatory mechanisms driving these transcriptional switches and their potential as therapeutic or transmission-blocking targets.

In summary, across its life cycle, *T. gondii* undergoes a stepwise metabolic reconfiguration tailored to each developmental niche. Tachyzoites rely heavily on lipid and branched-chain amino acid catabolism for rapid replication, while bradyzoites expand detoxification, redox buffering, and protective pathways for long-term persistence. Upon entering the feline intestine (EES1 → EES2), the parasite broadens its carbon and sulfur usage and recruits “specialized” routes. During gametogenesis (EES5), fatty acid and short-chain carboxylate metabolism intensify, and in oocyst stages, these themes persist and culminate in extensive secondary metabolism that fortifies fully sporulated oocysts for survival outside the host.

## Conclusions

This study establishes a comprehensive transcriptomic framework of the entire *T. gondii* life cycle, with particular emphasis on early timepoints in oocyst sporulation. By integrating newly generated RNA-Seq datasets of unsporulated, sporulating, and fully sporulated oocysts (endpoint) to previously published transcriptomes from tachyzoites, bradyzoites, and enteroepithelial stages, we close a critical knowledge gap and present the first genome-wide dataset covering all developmental phases of this parasite.

Our analyses demonstrate that sporulation in coccidia is accompanied by surprisingly extensive transcriptional reprogramming involving thousands of genes, while more limited adjustments characterize transitions within closely related stages occupying similar niches. These data provide clear evidence that developmental regulation in *T. gondii* is highly modular, with stage-specific transcriptional signatures that can be linked to defined biological functions. In particular, the prominent involvement of AP2 family transcription factors and other DNA-binding proteins underscores their importance in regulating key developmental transitions, including sporulation.

Metabolic pathway analysis revealed striking stage-specific rewiring. Tachyzoites preferentially activate pathways supporting rapid growth, while bradyzoites shift to detoxification and persistence. Within the highly specialized enterocyte niche in the feline intestine, carbon, sulfur, and amino acid metabolism expand to meet the rapid expansion during asexual development and the specific demands of gametogenesis. During oocyst maturation, the parasite engages an extensive network of secondary metabolic pathways, including terpenoid biosynthesis, menaquinone/ubiquinone metabolism, and specialized redox systems. In addition, we confirm the expression of protective factors such as LEA proteins and we show that most of the annotated OWPs are indeed expressed only during sporulation. Together, these expression patterns likely represent evolutionary conserved adaptive mechanisms to ensure survival and long-term environmental resilience of sporulated oocysts.

Importantly, our findings refine and expand upon earlier transcriptomic and proteomic studies, highlighting previously unrecognized gene sets and metabolic routes that may represent novel therapeutic or transmission-blocking targets. This may be particularly important for prioritizing future efforts to improve annotation of the *T. gondii* genome. By mapping expression profiles of oocyst wall proteins, sporocyst-associated factors, and stress-response genes, we also provide a valuable catalog of candidates for functional validation.

Taken together, this integrated transcriptomic atlas represents a major advance in the understanding of *T. gondii* development and gene expression trajectories, particularly the molecular events underpinning sporulation, ultimately informing strategies aimed at preventing parasite transmission and mitigating the global burden of toxoplasmosis.

## Data and Code Availability

RNA-Seq data have been deposited in GEO under accession number GSE206344. All R scripts used for variance filtering, clustering, differential expression analysis, ORA, and visualization are openly available at the UZH GitLab repository: [Hehl Lab-*Toxoplasma*_transcriptome_atlas](https://gitlab.uzh.ch/hehllab/toxoplasma_transcription_atlas_sporulation). Interactive UMAP visualizations are permanently hosted on RPubs at [High-variance 935 genes](https://rpubs.com/Elec/HighVariance935genesTGondii) and [Full transcriptome](https://rpubs.com/Elec/FullTranscriptomicsTgondii). The interactive network diagrams are permanently hosted on NDEX:([NDEX 1](https://www.ndexbio.org/viewer/networks/b977c071-88e0-11f0-a218-005056ae3c32), [NDEX 2](https://www.ndexbio.org/viewer/networks/b0998dd3-8a2d-11f0-a218-005056ae3c32), [NDEX 3](https://www.ndexbio.org/viewer/networks/a8c7164d-1611-11f0-9806-005056ae3c32), [NDEX 4](https://www.ndexbio.org/viewer/networks/92d45ead-8991-11f0-a218-005056ae3c32), [NDEX 5](https://www.ndexbio.org/viewer/networks/f1b8ba87-8980-11f0-a218-005056ae3c32)) for full interactivity and more details, open [Cytoscape](https://cytoscape.org/).

## Supporting information

Links to online resources

## Acknowledgements

We thank Prof. Nicholas C. Smith and Prof. Peter Deplazes for assistance with experimental infection of cats and purification of oocysts. We are grateful to Dr. Catharine Aquino, Dr. Timothy Sykes and the Functional Genomics Center Zurich for their support.

## Author information

### Additional information

#### Authors’ contributions

Conceptualization: ABH, CR; Data Curation: EM; Formal Analysis: ABH, EM, CR; Funding Acquisition: ABH; Methodology: CR, EM; Writing – Original Draft: ABH, CR, EM; Writing – Review & Editing: ABH, CR

