## Supplemental figure legends for "Genome-wide Transcriptomic Analysis of T*oxoplasma gondii* Reveals Stage-specific Regulatory Programs and Metabolic Adaptations Driving Oocyst Sporulation"

**Fig. S1**: Quality control of RNA and RNA-Seq data. **A** Percentage of sporulated oocysts (i.e. containing sporocysts) observed in each sample. **B** Simulated gel electrophoreses based on Agilent Tapestation using an RNA Screen Tape. The green lines mark the lower marker, red asterisks the parasite-specific 26S ribosomal RNA bands, green arrows the 18S rRNA bands. **C** Total and reads mapped to the *Toxoplasma gondii* ME49 genome (ToxoDB Release 68).

**Fig. S2**: Principal component analysis of all samples. Principal component analysis depicting all oocyst samples.

**Fig. S5**: Percentage of transcripts assigned to stages of their expression maxima identified by . by Fritz *et al.* [3]  (n = 165), **B** Possenti *et al.* [4] (n = 135)All IDs are corrected for ToxoDB Release 68 and only IDs from the reference genome ME49 have been considered. Non-existing IDs were not considered.
