## Supplementary figures and images for "Genome-wide Transcriptomic Analysis of T*oxoplasma gondii* Reveals Stage-specific Regulatory Programs and Metabolic Adaptations Driving Oocyst Sporulation"

### Supplemental figure 3

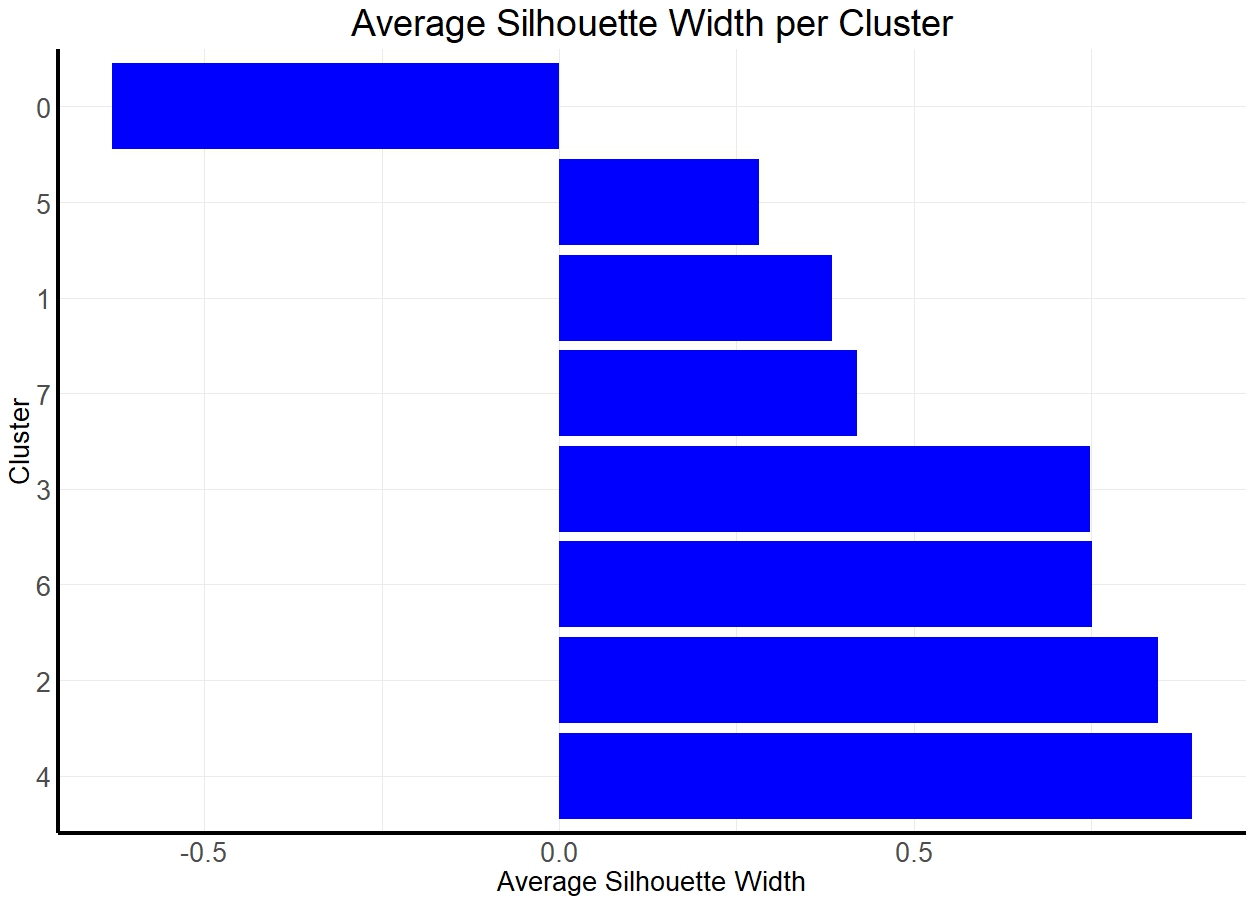

### Supplemental figure 4

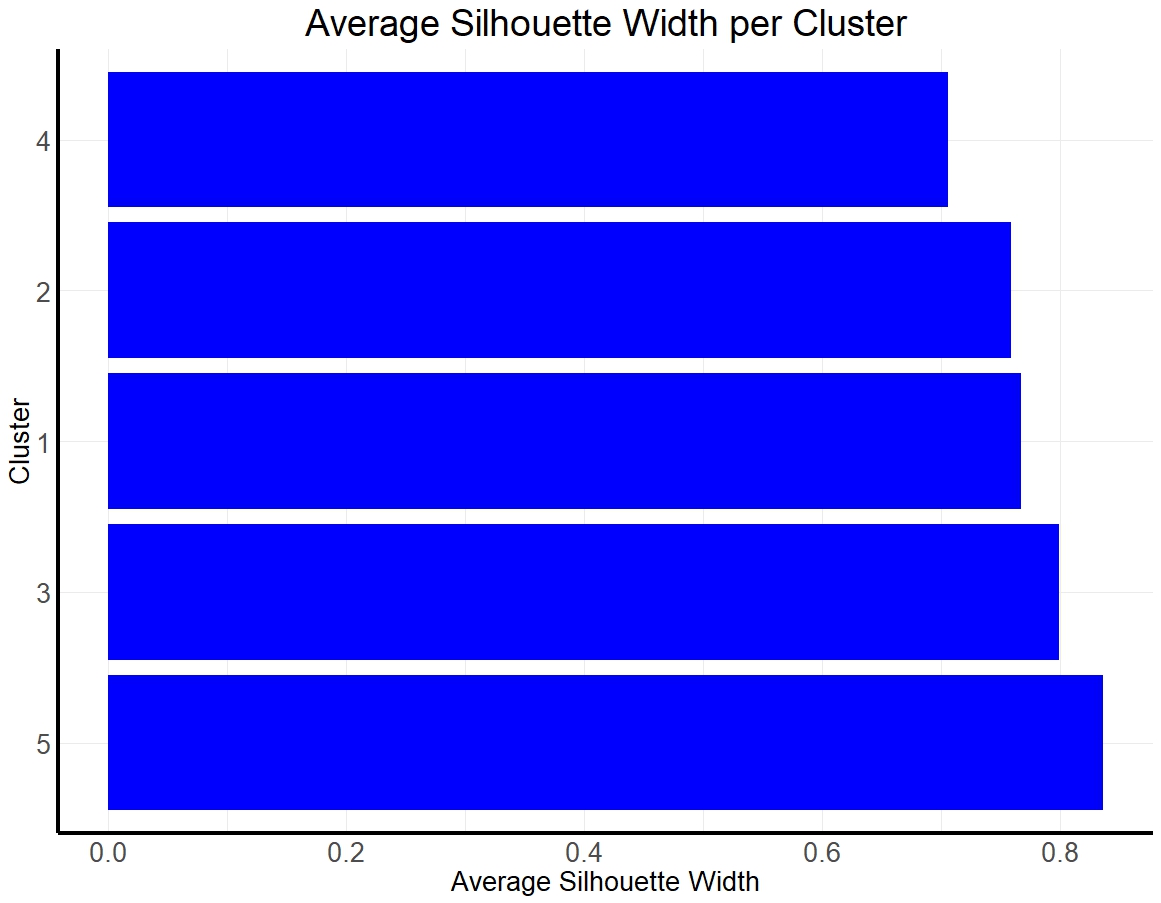

### Supplemental figure 5

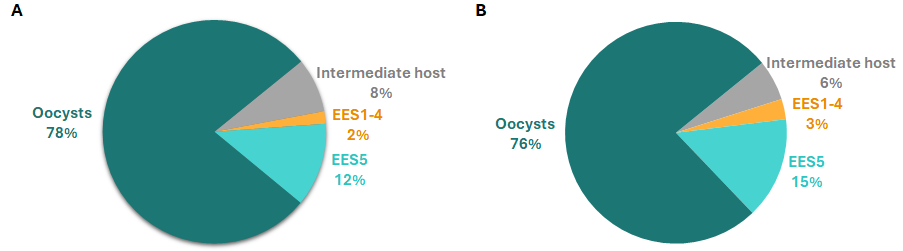

### Supplemental figure S1

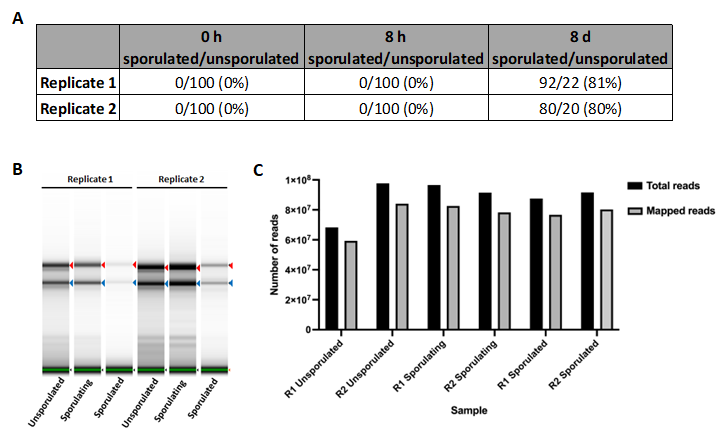

### Supplemental figure S2

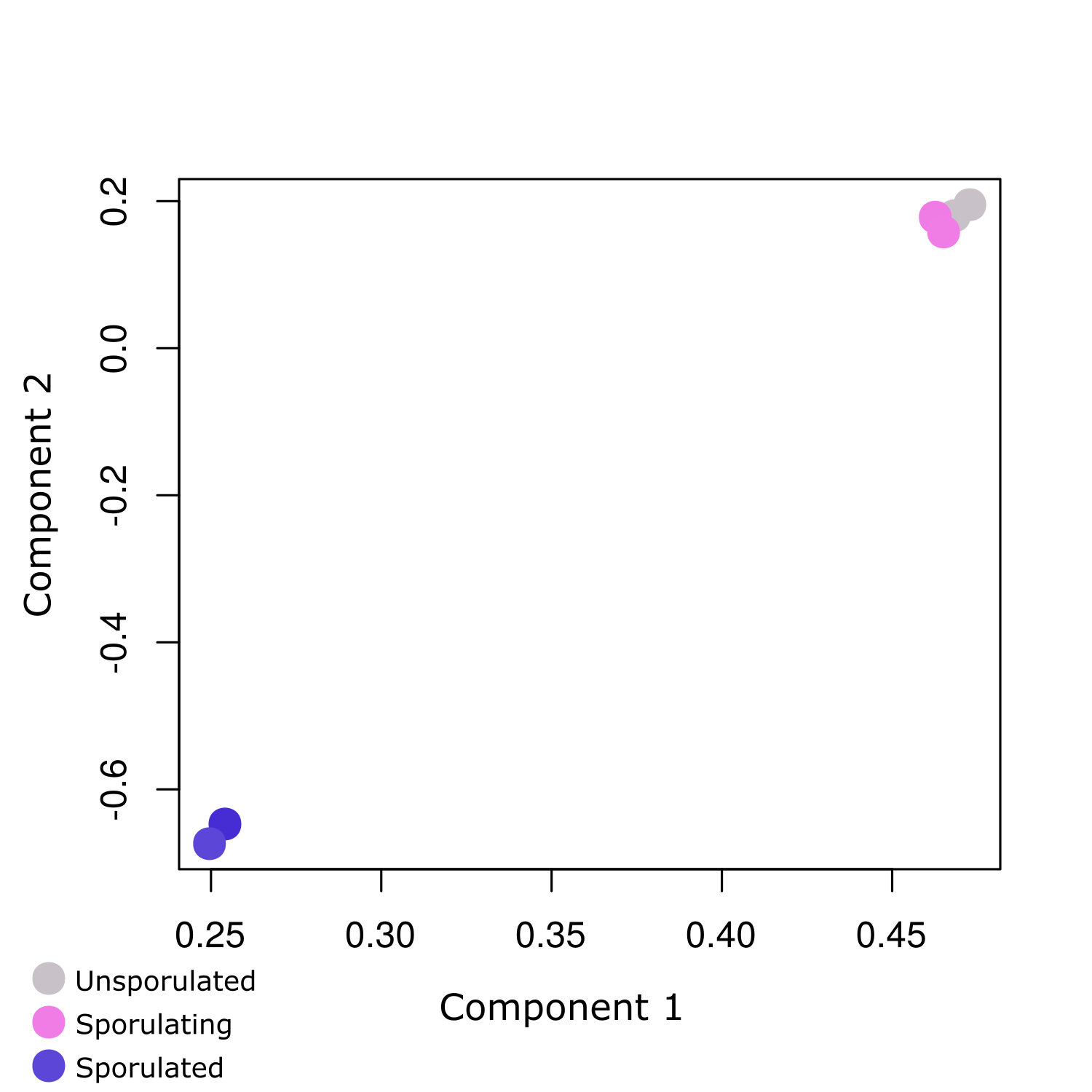
