## Supplementary material for "Genome-wide Transcriptomic Analysis of T*oxoplasma gondii* Reveals Stage-specific Regulatory Programs and Metabolic Adaptations Driving Oocyst Sporulation": Links to online resources

Magoye et al. 2025

**Links to online resources:**

<https://gitlab.uzh.ch/hehllab/toxoplasma_transcription_atlas_sporulation/-/tree/main/Data_Resources?ref_type=heads>

**1. R analysis scripts**

All scripts used for variance filtering, clustering,UMAP, DEA, ORA, and visualization are available at:
👉 [UZH GitLab Repository](https://gitlab.uzh.ch/hehllab/toxoplasma_transcription_atlas_sporulation)

**2. Transcriptome atlas (interactive UMAPs)**

- [RPubs: UMAP of Bulk RNA-seq (High-variance 935 genes)](https://rpubs.com/Elec/HighVariance935genesTGondii)
- [RPubs: UMAP of Bulk RNA-seq (Full transcriptome)](https://rpubs.com/Elec/FullTranscriptomicsTgondii)

**3. RNA-seq data**

Raw and processed RNA-seq data are deposited at GEO:
👉 [NCBI GEO Accession Viewer,Transcriptomics of T. gondii](https://www.ncbi.nlm.nih.gov/geo/query/acc.cgi?acc=GSE206344) *(Accession number:GSE206344)*

**4. Interactive ORA networks (NDEx)**

- [Tachyzoite → Bradyzoite transition](https://www.ndexbio.org/viewer/networks/b977c071-88e0-11f0-a218-005056ae3c32)
- [Bradyzoite → EES1 transition](https://www.ndexbio.org/viewer/networks/b0998dd3-8a2d-11f0-a218-005056ae3c32)
- [EES1 → EES2 transition](https://www.ndexbio.org/viewer/networks/a8c7164d-1611-11f0-9806-005056ae3c32)
- [EES5 → Unsporulated oocyst transition](https://www.ndexbio.org/viewer/networks/92d45ead-8991-11f0-a218-005056ae3c32)
- [Sporulating → Sporulated oocyst transition](https://www.ndexbio.org/viewer/networks/f1b8ba87-8980-11f0-a218-005056ae3c32)

**Notes**

- This directory contains **only metadata and links**.
- All analyses were performed in R (v4.4.1) with scripts provided in the main repository.
- For questions, please contact the **Hehl Lab,University of Zurich. (****)**.
